# HDCluster: High-Degree Graph Clustering for Robust Analysis of Single Molecule Localization Microscopy

**DOI:** 10.1101/2025.10.23.684134

**Authors:** Ismail M. Khater, Seyed Benyamin Bagheri, Shane J. Eastwood, Ivan Robert Nabi, Ghassan Hamarneh

**Affiliations:** Department of Electrical and Computer Engineering, Faculty of Engineering and Technology, Birzeit University, Birzeit P627, Palestine; Medical Image Analysis Lab, School of Computing Science, Simon Fraser University, Burnaby, BC V5A 1S6, Canada; Department of Genetics, Harvard Medical School, Boston, MA, 02115, USA; Life Sciences Institute, Department of Cellular and Physiological Sciences, University of British Columbia, Vancouver, BC V6T 1Z3, Canada; School of Biomedical Engineering, University of British Columbia, Vancouver, BC V6T 1Z3, Canada

## Abstract

Clustering is a fundamental task in data analysis: grouping similar objects together into distinguishable subsets. Here, we introduce HDCluster, a novel high-degree graph-based clustering algorithm designed to effectively and rapidly handle various real-world clustering applications, particularly in the context of super-resolution single molecule localization microscopy (SMLM). HDCluster efficiently handles datasets with large and variable numbers of clusters, without requiring prior knowledge of the cluster count, relying on only one parameter. The high speed and efficiency of HDCluster allow it to handle large SMLM datasets with millions of localizations. A comprehensive quantitative comparison shows that HDCluster is equivalent to state-of-the-art clustering methods based on clustering performance measures, such as ARI and AMI, on simulated datasets. In terms of time efficiency, HDCluster significantly outperforms other clustering algorithms on large DNA-PAINT datasets. Application of HDCluster to a neuronal DNA-PAINT dataset on synapse biology identified and removed multiple blinking artifacts and effectively identified excitatory and inhibitory synapses, highlighting its utility for real experimental DNA-PAINT data. HDCluster is particularly robust to noise, making it a promising and effective tool for various clustering tasks in big-data settings, such as SMLM DNA-PAINT datasets.

## INTRODUCTION

Clustering is an unsupervised machine learning task where data samples are grouped according to similarities, ensuring that similar samples fall into the same group while dissimilar ones are separated. Hence, grouping is formed based on intrinsic similarity patterns across the data^1^. Data clustering is valuable in domains where ground truth labels are not available or hard to collect, such as single molecule localization microscopy (SMLM).

SMLM comprises a family of super-resolution imaging techniques, such as Photoactivated Localization Microscopy (PALM)^2^ and Stochastic Optical Reconstruction Microscopy (STORM)^3^, that circumvent the Abbe diffraction limit of conventional light microscopy via stochastic excitation of fluorophores and fitting of the point spread function (PSF) to localize single fluorophores at precisions of tens of nanometers^4^. DNA points accumulation for imaging in nanoscale topography (DNA-PAINT) utilizes the transient binding of fluorescent DNA strands to achieve both high precision and multiplexing^5^. Minimal photon fluxes (MINFLUX) combines spatially structured illumination with localization analysis^6^ and Resolution Enhancement by Sequential Imaging (RESI) combines DNA-PAINT with sequential imaging of sparse labeling sites^7^ achieving nanometer and sub-nanometer resolution. SMLM data are represented as a point cloud of 2D/3D coordinates of molecular localizations^8^. SMLM data cluster analysis enables quantitative study of nucleic acid organization, protein aggregates, and other cellular structures at the nanoscale^9–11^. However, SMLM data exhibit challenges such as noise, variations in density and size, large dataset volumes, and the presence of a high and often unknown number of clusters^12^.

Several algorithms have been proposed for data clustering in general, and specifically for SMLM data analysis, based on diverse principles, including distance, density, graph/spectral properties, or other similarity-driven approaches^12^. Every clustering algorithm has underlying assumptions and parameters that restrict its universality across applications and data types. For example, the k-means algorithm requires the number of clusters to be specified ahead of time and assumes spherical clusters of similar sizes^1,13^. Other clustering methods are sensitive to noise and outliers (e.g., hierarchical clustering), lack scalability to very large datasets in terms of time and space (i.e., memory), and struggle with clusters of varying shape and density^1,13^. Here, we introduce HDCluster, a similarity graph-based clustering algorithm, and show that it outperforms competing clustering algorithms and is particularly robust to the challenges of noisy, heterogeneous point cloud-based SMLM DNA-PAINT data. HDCluster supports various computational tasks aimed at SMLM data analysis, including reconstructing binding sites, identifying nanoclusters and macromolecular structures, and denoising the data. Application to real DNA-PAINT datasets shows that HDCluster exhibits significantly improved time efficiency and effectively removes multiple blinking artifacts.

## RESULTS

### 1. HDCluster Framework and Computational Tasks

HDCluster is a spatial data clustering method that can be leveraged to find clusters in SMLM data. The HDCluster algorithm has one parameter, merging threshold (*mT*ℎ), used to construct the similarity graph and get the node degree as a feature to start iterating and merging the nodes to get the clusters. The setting of the *mT*ℎ parameter is related to the physical characteristics of the clusters, i.e., the size/scale of the sought clusters (Figure 1A). HDCluster finds the clusters after the convergence condition is achieved and returns the per-localization class label, including labels for noisy localizations. β is an optional parameter to control noise removal with a default setting that can be changed if needed. See the Methods section for more details about the algorithm.

**Figure 1.**
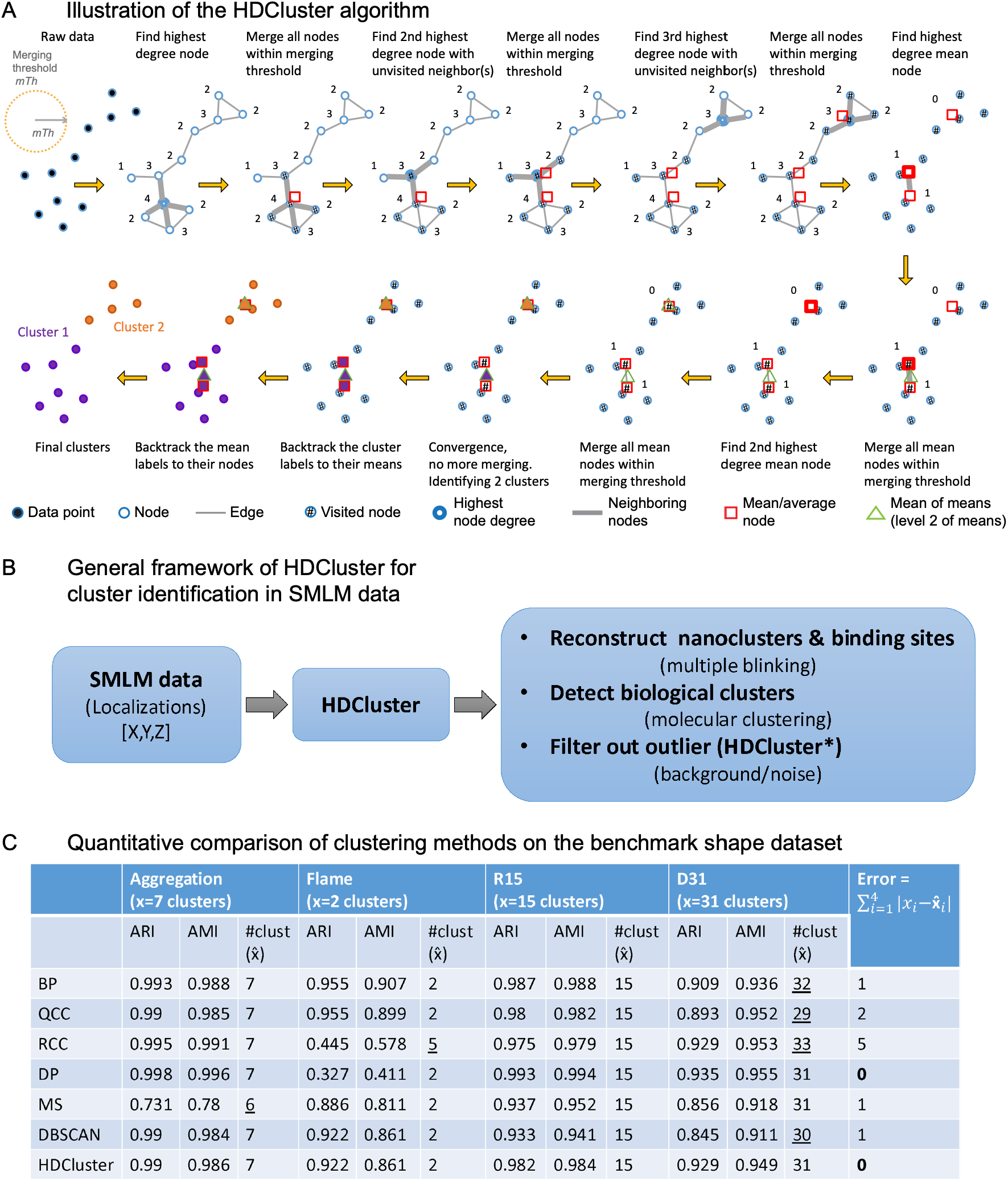
Overview of the HDCluster framework to cluster, denoise, and reconstruct the emitters/binding sites in simulated and DNA-PAINT data. (A) Illustration of the HDCluster algorithm to obtain data clusters as well as the centroids (the consensus localizations that approximate the true emitter positions by computing the mean of means) of the extracted clusters. More formally, HDCluster pseudocode is provided in Algorithm 1. HDCluster is an iterative algorithm that starts from the similarity graph constructed based on the *mT*ℎ parameter. The degrees of the graph nodes are calculated. The nodes are sorted in descending order of node degree. The algorithm starts iterating from the node with the highest degree. All the nodes that share edges with this high-degree node are merged; the mean (in terms of average coordinates for the nodes’ spatial locations) of these merged nodes is calculated and all these merged nodes are marked as visited nodes. After that, the algorithm moves to the next highest degree node in the list, with at least one node that is marked as not visited, and so on until all the nodes are marked as visited. After concluding this first round of iterations, HDCluster enters another round by constructing an updated graph based on the mean nodes that were produced from the node merging of the first round of iterations. The algorithm will start iterating again from the mean node with the highest degree, as it did in the first round, and keep iterating and producing the mean of means until the convergence condition is reached. The convergence condition is reached when the mean nodes have no edges (i.e., the degree of the top-level mean nodes equals zero). Every one of the high-level mean nodes will be considered as a top-level cluster ID and will be used to backtrack the cluster labels to their means and so on down to the nodes that the means come from. The algorithm keeps propagating/backtracking the labels from the top-level mean node down to the nodes, and all the nodes will get an ID or label based on the top-level mean node. The algorithm does so for the rest of the top-level mean nodes. At the end, all the nodes get labeled and have membership in one and only one cluster with a unique ID. The top-level mean nodes are the emitters (consensus localizations that approximate the true emitter positions) of the nanoclusters in SMLM data. HDCluster is capable of filtering out the noisy localizations in addition to data clustering. (B) A general framework of SMLM data analysis and the computational tasks that HDCluster can perform for SMLM data analysis. (C) Quantitative analysis (ARI, AMI, #clusters) of clustering methods evaluated on the benchmark shape dataset^19^ presented in Supp. Fig. 1. The best parameter(s) for each method are selected based on the AMI measure. See Supplementary Videos for an animation of the HDCluster algorithm workflow and its sensitivity to parameter adjustments.

Figure 1B illustrates HDCluster capabilities, encompassing various computational tasks for SMLM data analysis, such as (1) emitter reconstruction (or binding sites reconstruction^14^), (2) finding true biological clusters and nanoclusters in the SMLM data, and (3) denoising SMLM data.

### 2. HDCluster Performance Comparison with Other Clustering Methods

To evaluate clustering performance and tune the algorithm’s parameters, we used the adjusted Rand index (ARI)^15^ and the adjusted mutual information (AMI)^16^ as performance metrics. ARI measures similarity between predicted and ground truth labels and AMI captures the amount of shared information between clusters, corrected for chance. AMI is a more robust measure when the number of clusters varies and accounts for bias toward certain cluster size distributions^17,18^.

We applied HDCluster and other commonly used clustering methods to the shape benchmarking dataset^19^, widely used for assessing clustering methods’ performance. This dataset of 2D points and their corresponding membership as ground truth (GT) labels includes blob-like and spherical clusters similar to SMLM data of biological clusters: Aggregation with 7 clusters, Flame with 2 clusters, R15 with 15 clusters, and D31 with 31 clusters (Supp. Figure 1A). We tested state-of-the-art clustering methods, including Border Peeling (BP)^20^, Quasi-Cluster Centers (QCC)^21^, Robust Continuous Clustering (RCC)^22^, Density Peaks (DP)^23^, mean-shift (MS)^24^, and Density-Based Spatial Clustering of Applications with Noise (DBSCAN)^25^, as well as HDCluster, on these benchmark datasets (Supp. Table 1 shows the number of parameters and other information for the studied methods). We tuned each method’s parameters based on AMI. Supp. Figure 1 shows the qualitative results and Figure 1C summarizes the quantitative results when applying the clustering methods to the shape benchmarking dataset. Only HDCluster and DP retrieve the correct number of clusters across all tested datasets. However, DP shows poor performance on the Flame dataset (ARI of 33% and AMI of 41%), where HDCluster achieves much higher accuracy (92% ARI and 86% AMI). Notably, HDCluster outperforms DBSCAN on the D31 and R15 datasets, both of which mimic nanocluster structures observed in SMLM data. The benchmark shape datasets demonstrate that HDCluster provides consistently strong performance across diverse clustering scenarios. While other methods may achieve higher scores on specific datasets, HDCluster maintains competitive clustering accuracy and is among the few methods that correctly identify the expected number of clusters across all benchmark datasets. These results highlight the robustness of HDCluster to variations in cluster shape, density, and spatial arrangement.

Graph-based clustering methods such as QCC and RCC construct similarity graphs from the data; where QCC via KNN/reverse-KNN density estimation, and RCC via a k-NN-based connectivity structure combined with global continuous optimization. In our experiments (Supp. Table 1), both approaches showed limited scalability to datasets on the order of tens to hundreds of thousands of points, consistent with the increased computational and memory demands of maintaining these graph structures and, for RCC, its optimization procedure. In contrast, HDCluster constructs a sparse ε-graph (i.e., a radius-based graph) using KD-tree neighbour search and performs degree-guided merging, thereby progressively reducing the number of representatives without requiring global optimization, enabling efficient clustering of millions of points on standard hardware.

Nieves et al.^26^ introduced a standard benchmarking framework for comparing clustering algorithms on SMLM point cloud data. Their work includes generating simulated ground truth point patterns with realistic blinking and localization uncertainty. The dataset comprises 10 distinct scenarios/conditions (variations in density, cluster size, blinking behavior, background noise, etc.) with and without multiple blinking artifacts. Each scenario contains 50 different point cloud files. They tested and compared the performance of several clustering methods, which include: DBSCAN^25^, topological mode analysis tool (ToMATo)^27^, kernel density estimation (KDE)^28^, fast optimized clustering algorithm for localizations (FOCAL)^29^, cluster analysis by machine learning (CAML)^30^, ClusterViSu^31^, and SR-Tesseler^32^. The methods’ parameter spaces are scanned systematically under each condition. We assessed HDCluster using the benchmarking SMLM datasets in accordance with the methodology outlined by Nieves et al.^26^. All experiments were performed with the denoising feature activated. We tuned the key parameters of HDCluster, *mT*ℎ and the optional filtering parameter β, to maximize the average AMI of the 50 files of each scenario.

Supp. Figure 2 shows HDCluster’s ARI results, a collection of Nieves et al.’s results^26^, and additional data obtained by Pineda et al.^18^, with Scenarios 2-10 being compared across the various methods mentioned on the benchmark dataset. Supp. Figure 2A shows results on the ground truth scenarios of the benchmark, while Supp. Figure 2B shows the results of multiple blinking scenarios. We observed that Pineda et al.^18^ had values that differed from Nieves et al.^26^, and we decided to use the optimal parameters set by Nieves et al.^26^, and collect new ARI scores. In Nieves et al.’s results, DBSCAN, ToMATo, and KDE generally perform well across multiple conditions and are relatively robust, with DBSCAN showing superiority. The other methods, ClusterViSu, SR-Tesseler, and FOCAL, show weaker performance in several settings. All methods suffer performance drops under multiple blinking localization conditions. Our DBSCAN analysis, despite using the same optimal parameters, performed lower than reported in Nieves et al.^26^. Of note, DBSCAN as implemented by Pineda et al.^18^ outperformed DBSCAN as implemented by either ourselves or Nieves et al.^26^, and matched the performance obtained with MIRO^18^ (Pineda et al.’s method) (Supp. Figure 2). Discrepancies between performance values for DBSCAN may relate to how DBSCAN’s algorithm handles the border points, which becomes more prevalent when implemented on different workstations^33^. Recent SMLM clustering approaches have explored application of machine learning and deep learning techniques (e.g., MIRO^18^) to address challenges such as noise, heterogeneity, and emitter separability. While these methods show promising performance (Supp. Figure 2), they typically require labeled training data, which is often unavailable or difficult to obtain in experimental SMLM settings. In contrast, HDCluster is an unsupervised, training-free approach, which is an important strength when there is an absence of labeled training data, such as required for the performance gains from MIRO.

For performance comparison, we selected DBSCAN, the most effective method identified by Nieves et al.^26^, and employed their reported parameters to ensure fairness. The AMI and ARI values for HDCluster and DBSCAN are presented in Supp. Table 2. Across all simulated scenarios, HDCluster matches DBSCAN in both ideal (i.e., no added blinking) and multiple blinking scenarios. In the ideal scenarios, both methods achieve comparable results, with HDCluster showing slightly higher ARI and AMI values in most cases (e.g., Scenario 3: 0.90 ± 0.017 vs. 0.84 ± 0.017 for ARI). However, when multiple blinking is introduced, both methods show mixed behavior. HDCluster achieves slightly higher AMI values on average (0.681 ± 0.040) compared to DBSCAN (0.676 ± 0.043), indicating comparable cluster agreement. Yet, DBSCAN achieves slightly higher ARI values (0.577 ± 0.048) than HDCluster (0.570 ± 0.046). These differences are small, suggesting similar clustering performance under the more challenging multiple blinking conditions. All results reported in Supp. Table 2 were generated independently using our implementation and evaluation pipeline. For DBSCAN, the algorithm was executed using the optimal parameter values (ε and MinPts) reported by Nieves et al.^26^, while the performance metrics were recalculated from the resulting cluster assignments using our evaluation framework.

Overall, the benchmarking results indicate that HDCluster maintains clustering performance comparable to that of DBSCAN in both ideal and multiple blinking scenarios. While neither method consistently outperforms the other across all metrics and scenarios, HDCluster achieves similar clustering accuracy while offering additional practical advantages, including a simpler parameterization, robust performance across diverse datasets, and substantially lower computational cost (see Supp. Table 4). These findings suggest that HDCluster provides an effective and computationally efficient alternative to DBSCAN for clustering SMLM point-cloud data, including datasets affected by multiple blinking artifacts.

For fair and direct comparison with the benchmarking framework of Nieves et al.^26^, we evaluated clustering performance using ARI and AMI, which are widely adopted metrics in SMLM clustering studies. While these measures quantify agreement with ground-truth partitions, they do not fully capture additional properties such as localization precision, cluster compactness, or emitter separability under realistic blinking conditions. Recent work by Pineda et al.^18^ has proposed complementary evaluation metrics tailored to SMLM data (see also Supp. Figure 5 and Equations 3–5 for measures newly introduced in this work). Incorporating such measures could provide a more comprehensive assessment and will be considered in future work.

### 3. HDCluster Robustness to Noise and Scalability

To test the ability of the different clustering methods to handle noisy data, we generated datasets that comprise clusters with various shapes and densities with additive noise points as described in Methods Section 5. Figure 2A shows one experiment for the generated clusters concatenated with a noise level of 20% of the total clustered points drawn from a uniform distribution. DBSCAN and HDCluster efficiently handle the noisy data and extract clusters, while QCC, DP, RCC and MS are unable to accurately segment noise from clusters. We varied the noise level from 0% (clusters only) to 100% with a step size of 10% and compared DBSCAN, BP and HDCluster at each noise level (Figure 2B), where noise level is defined as the percentage of noise points relative to the total number of clustered (non-noise) points. Specifically, we simulated 3D datasets comprising 80 Gaussian clusters randomly distributed within a cubic volume spanning 0–1000 nm along each spatial dimension. To emulate clusters with varying densities, the number of points per cluster was randomly sampled between 10 and 100. Cluster shapes were generated as ellipsoidal Gaussian distributions with varying anisotropy, where the diagonal variances of the covariance matrix were independently sampled from a uniform distribution in the range of 1–21 nm, as described in Equation 7. Noise points were generated independently from a uniform spatial distribution across the entire field of view and were not restricted from overlapping with true clusters. Consequently, some noise points may, by chance, fall within or near cluster boundaries, thereby creating realistic and challenging clustering scenarios. The parameter(s) for DBSCAN and HDCluster are selected based on ground truth data and the AMI score, as shown in Figure 2C, which shows a single case where the noise level is 100%. DBSCAN and HDCluster are robust to high noise levels; for BP, AMI and ARI are progressively reduced as the noise level exceeds 40% (Figure 2B). Furthermore, the default value of β=0.2 was sufficient to effectively remove noise across a wide range of simulated noise levels (Figure 2B), indicating that HDCluster is not highly sensitive to choice of β.

**Figure 2.**
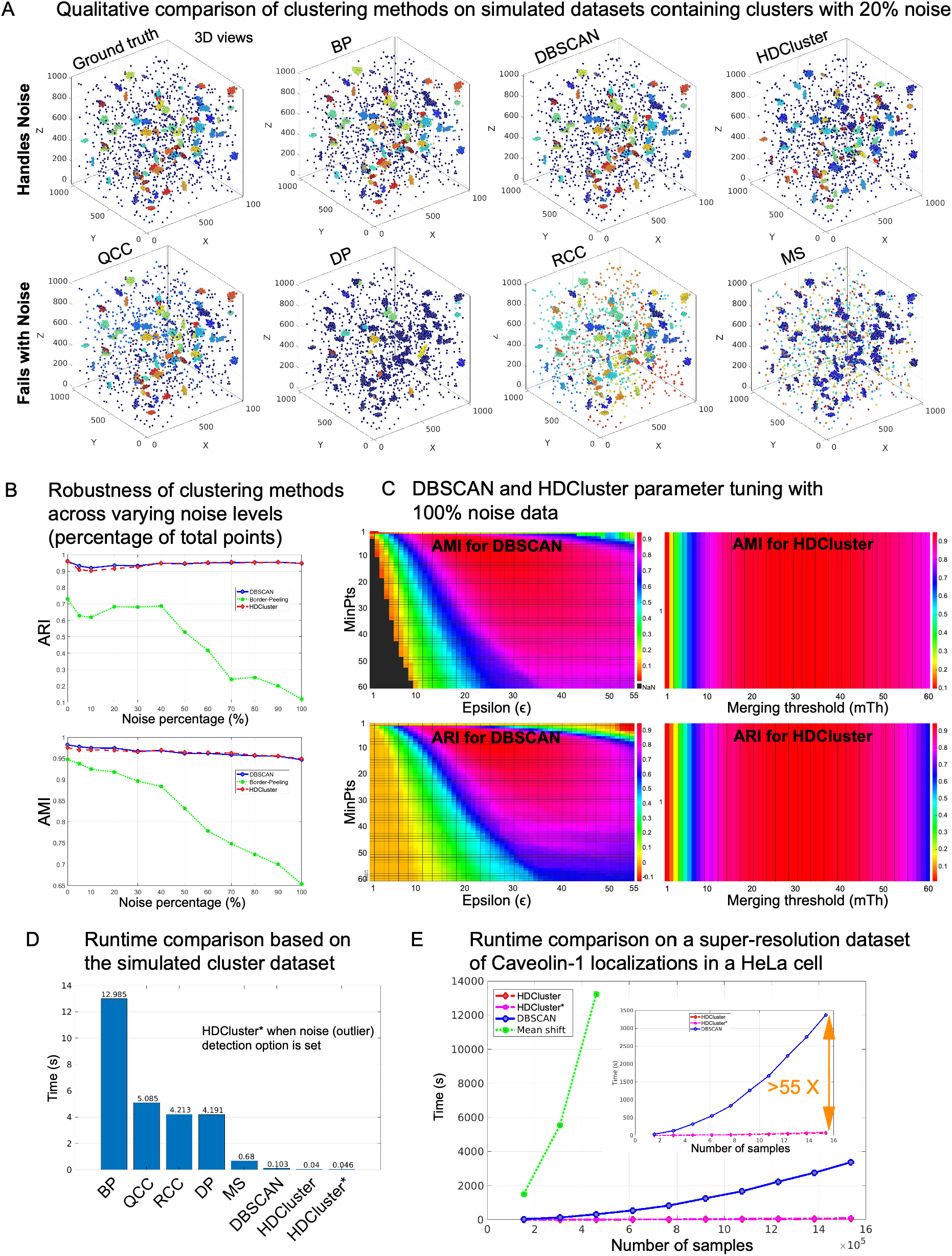
Noise robustness analysis of various clustering methods at different noise levels. Runtime comparison of the clustering methods based on the simulated and real SMLM dataset. (A) Simulated 3D data of 80 Gaussian clusters with various densities and ellipsoidal shapes is generated with various noise levels. In this simulation, we show the data with a 20% noise level of the total number of localizations. The various clustering methods have been applied to find the clusters and label the noisy localizations. For the parametric methods, the method’s parameter(s) are tuned based on the AMI. (B) Comparison of the clustering methods with the denoising capability for various noise levels. Some methods can handle noise but are not robust to high noise values in the data. (C) Parameter selection for the DBSCAN and HDCluster methods when applied to data with a 100% noise level. Both ARI and AMI measures were plotted for both methods at various parameters. The parameters with high AMI value were selected to report the results presented in (B). (D) Calibrate the runtime for the various clustering methods when applied to the simulated dataset presented in (A). All the methods were run on the same computer and the same dataset for fair comparison. Not all the methods can be used on large datasets. (E) Runtime analysis for the methods that can be run on large datasets, such as SMLM data. HDCluster, DBSCAN, and mean-shift (MS) algorithms were run on a dataset of Caveolin-1 localizations^34^ presented in the Supp. Figure 3. We start running the clustering methods on 10% of the whole data and calibrate the runtime, then we increase the localizations by 10% until we reach 100% of the total localizations. Not all the methods scale with the dataset size. HDCluster* refers to the version of the algorithm with the noise-handling option enabled.

To assess the speed of the clustering algorithm for SMLM data, we applied the clustering methods to the same datasets and on the same hardware (Intel(R) Core(TM) i7-4790 CPU @ 3.6GHz. Memory 32 GB, without other processes/applications running during testing). We first calibrated the running time for each algorithm on the small dataset from Figure 2A. Some of the algorithms are implemented in Python, while others are implemented in MATLAB. In our comparison, we used the latest optimized implementation of DBSCAN in MATLAB 2024b. HDCluster is the fastest algorithm, retrieving clusters in ∼0.046 seconds, with DBSCAN second followed by MS, DP, RCC, and QCC, and finally the BP algorithm (Figure 2D). We then applied the algorithms to a real SMLM dataset of caveolin-1 labeled HeLa cells of more than 1.5 million localizations^34^ (Supp. Figure 3). Only HDCluster, DBSCAN and MS were able to handle the data on our testing workstation (Intel(R) Xeon(R) CPU E5-2630 v3 @ 2.40GHz. Memory 94 GB) with the other algorithms crashing the computer and failing to complete. We calculated the runtime for MS, DBSCAN, and HDCluster starting with 10% of total localizations, progressively increasing the sample size to the total number of localizations (Figure 2E). The runtime for MS grew exponentially with dataset size, and we stopped the analysis after reaching 30% of the 1.5 million localizations. For DBSCAN and HDCluster, runtimes were calculated for increments up to the full data size (i.e., 100%, ∼1.5 million localizations). At 100% of the total data, HDCluster is >55 times faster than DBSCAN, and we expect the trend to continue as the dataset becomes larger (Figure 2E).

Supp. Table 4 expands on this further with the inclusion of Python implementations and comparison of an additional LifeAct actin dataset^35^. Local computer runtimes were computed on a workstation with AMD Ryzen 9 7900X @ 4.7GHz and 64 GB of DDR5 memory using the optimal parameters stated in Supp. Table 3. Scenarios 4 and 9 were chosen for comparison with the results provided by Nieves et al.^26^, as they comprise the scenario with the smallest quantity and lowest density of localizations (Scenario 4) and the one with the largest quantity and density of localizations (Scenario 9). Implementations of both DBSCAN and HDCluster in both MATLAB and Python were done for comparison, with HeLa caveolin-1 and LifeAct actin datasets used to show the impact of real scenario increased number of localizations and density on both algorithms. The Python implementation was done using scikit-learn’s DBSCAN with n_jobs = −1 for multi-threaded performance. An additional DBSCAN method by Wang et al^36^ was also applied to observe its speed impact. Both Scenarios 4 and 9 show strong preference for Python for the dataset initially, and also show the speed benefits of DBSCAN in Python, particularly the recent development of a parallel method which performs very fast on small datasets^36^.

For larger datasets, such as HeLa caveolin-1 (∼1.5 million localizations), DBSCAN in MATLAB proceeds much slower due to the implementation requiring to visit every point and recompute its neighbourhood. The algorithm iteratively performs an ɛ-neighbourhood search for each point encountered during cluster expansion, with MathWorks noting that distances are computed at each point of the iteration unless pairwise distances are precomputed^37^. HDCluster’s MATLAB implementation was much faster, as observed on another system (Figure 2E). Python implementations of both HDCluster and DBSCAN are slower than HDCluster MATLAB, with HDCluster Python being slower still. However, the parallel DBSCAN implementation^36^ showed incredibly rapid performance on this dataset. Analysis of DNA-PAINT datasets (LifeAct actin labeling, ∼65 million localizations)^35^ highlights some practical limitations of DBSCAN, especially when considering the parallel implementation^36^. The parallel DBSCAN implementation^36^ failed on the local workstation due to the large amount of memory required to process the dataset. Extra memory was also needed for the parallel DBSCAN versions in order to run to completion. In contrast, HDCluster effectively completed processing this large DNA-PAINT dataset with runtimes in Python similar to that of parallel DBSCAN processed on Digital Research Alliance of Canada cloud compute cluster. For comparison, HDCluster was also run on the same cluster, which ran slower, but this is due to the lower CPU frequency of the server processor. It is worth noting that the scikit-learn DBSCAN took twice as long on both local and the cloud cluster compared to HDCluster for the LifeAct actin dataset. Future parallelization of HDCluster may further speed up the algorithm, as observed for parallel DBSCAN^36^.

For MATLAB, due to the DBSCAN implementation, the run did not finish after over 50 hours of runtime. Although precomputing pairwise distances can improve runtime, this approach is infeasible for datasets containing tens of millions of localizations because the required distance matrix scales quadratically with the number of observations and the amount of memory required would be substantial.

### 4. HDCluster Binding Site Reconstruction Outperforms DBSCAN

Beyond cluster identification, binding site (emitter) reconstruction from repeated localizations is a pivotal task in SMLM data analysis (Figure 1B) that enables downstream applications such as protein counting, oligomerization analysis, and nearest-neighbour-based structural characterization^14,38^. To test whether the representative (or consensus) localizations of HDCluster iterative merging approximate true emitter positions, we analyzed SMLM data from the Gatta-PAINT 80 nm nanoruler that has three binding sites (emitters) organized linearly with a spacing of 80 nm (Supp. Figure 4A). Calculation of the 1st–5th nearest neighbour (NN) distances calculated for HDCluster reconstructed emitters shows that the majority of emitters are within ∼80 nm, as depicted in the first peak of the NN distances histogram (Supp. Figure 4A), while the rest of the emitters are within ∼160 nm (the nanoruler extreme emitter dimensions) as depicted in the second peak of the same histogram. As expected, HDCluster not only recovered the actual position of the emitters of the nanoruler rod but also preserved the relative distances and recovered the actual geometry of the designed nano-structure. HDCluster was then applied to the DNA origami nanostructure dataset published recently for the SPINNA method that compares experimental single-protein NN distances with simulated data generated from user-defined protein oligomerization models^38^. HDCluster was used to reconstruct the binding sites for linear and triangular DNA origami nanostructures with three binding sites spaced ∼15 nm (Supp. Figure 4B). We used *mT*ℎ = 0.04 × *pixelSize* (i.e., *mT*ℎ = 5.2 *nm*), where the pixel size is 130 nm. For linear origami, HDCluster effectively reconstructs the binding sites for the two origami structures and calculates NN distances, where the first peak is at ∼15 nm and a second peak at ∼30 nm for linear origami and ∼15 nm for triangular nanostructures, which agrees with the true dimensions and geometry of the designed structures.

We then compared the binding site reconstruction performance of HDCluster and DBSCAN using a dataset of 3 × 4 origami grids arranged at 20 nm spacing (Figure 3A)^39^. HDCluster was applied with *mT*ℎ = 0.06 × *pixelSize* (i.e., *mT*ℎ = 7.8 *nm* for a pixel size of 130 nm) and evaluated against the results reported by Stein et al.^39^, who analyzed NN of DBSCAN cluster centers using the same parameters reported in their study: *ε* = 2 × *σ_NeNA_* = 2 × 4.2 = 8.4 *nm* and *MinPts* = 6. Calculations of the 1st–5th NN distance show that the peaks are sharper and clearer for HDCluster relative to DBSCAN, with the highest peak at 20 nm (peak (a) in Figure 3B), the second highest peak at 28 nm (peak (b)) and so on (Figure 3B). The zoomed-in regions show superior clustering performance when using HDCluster over DBSCAN. With DBSCAN, some of the binding sites are filtered out, some merged and some averaged, affecting counting and the subsequent analysis of NN values. Indeed, the frequency of occurrences is more representative for HDCluster than DBSCAN; the expected 17 occurrences of peak (a) vs 8 occurrences of peak (d) in the 3 × 4 origami grids are accurately reported by HDCluster but not DBSCAN, which shows a similar frequency of events for peaks (a) and (d). Overall, HDCluster outperforms DBSCAN in binding sites reconstruction and localization.

**Figure 3.**
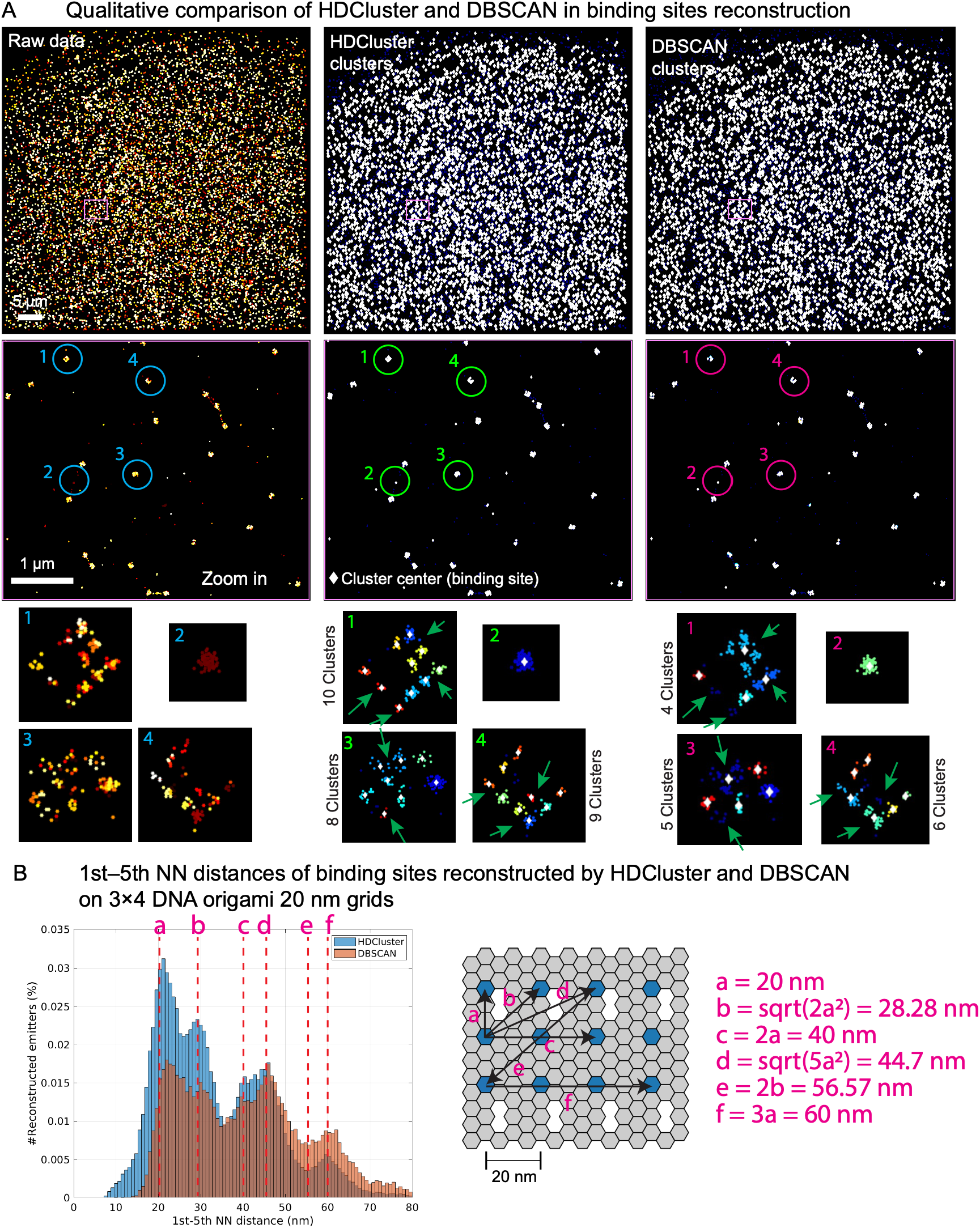
Qualitative and quantitative comparison of binding sites reconstruction and quality of clustering when using DBSCAN and HDCluster when applied to 3 × 4 DNA origami dataset^39^. (A) The quality of clusters impacts the binding sites’ subsequent analysis, as in the nearest neighbour (NN) analysis. We adopt the same DBSCAN parameters used in Stein et al.^39^ to find the clusters. DBSCAN can find the clusters, but when the clusters overlap, it might get them wrong. On the other hand, HDCluster can easily be adopted with one parameter to find more accurate clusters and consequently the binding sites. (B) Quantitative analysis of the reconstructed binding sites from both DBSCAN and HDCluster methods. The NN analysis for the binding sites from the 3 × 4 DNA 20 nm origami grids shows that the histogram peaks correspond to the geometrical grid of sites that distant 20 nm apart for both DBSCAN and HDCluster methods. However, the HDCluster is more realistic, while DBSCAN has limitations for the overlapped clusters and is sensitive to noisy points.

### 5. Application to Real DNA-PAINT Datasets

To test the utility of HDCluster for real DNA-PAINT datasets, we applied it to a published 13-plex neuronal labeling experiment incorporating cytoskeletal, organelle and synaptic markers^35^. To define the appropriate *mT*ℎ to cluster local multiple blinking localization networks, we used Ripley’s H function to identify the minimal cluster size. Ripley’s H analysis identified a first small-scale deflection from the initial slope near 6 nm across all 13 channels (Figure 4A). The consistency of this scale across channels suggests that it is related to features common to the multiplexed DNA-PAINT labeling protocol rather than channel-specific biological organization^35^. To test the impact of *mT*ℎ, we plotted the number of detected clusters at each *mT*ℎ relative to the number detected at *mT*ℎ = 1 nm for each of the six synaptic markers (Figure 4B). The initial increase in normalized cluster number relative to *mT*ℎ = 1 nm was observed at low *mT*ℎ values. Importantly, the percent change in centroid number decreased and stabilized around *mT*ℎ = 6 nm. We further assessed the effect of the optional denoising parameter β and found that centroid recovery was insensitive to β up to approximately the Ripley-derived 6-nm scale, with greater β-dependent differences at larger *mT*ℎ values (Figure 4C; Supp. Figure 6). Insensitivity of centroid recovery to β suggests that, at *mT*ℎ values up to the Ripley-derived scale, denoising effectively removes isolated zero-degree localizations (see Equation (6) and Algorithm 2 line 3). This is consistent with oversampled, noisy DNA-PAINT data containing local multi-blinking clusters of localizations as well as an equal or larger number of isolated, at the applied *mT*ℎ, zero-degree localizations. While further analysis of diverse DNA-PAINT datasets is required, our analysis suggests that for real, noisy DNA-PAINT data, given the observed insensitivity to β, HDCluster can be implemented as a one-parameter clustering algorithm with *mT*ℎ determined from Ripley’s H analysis.

**Figure 4.**
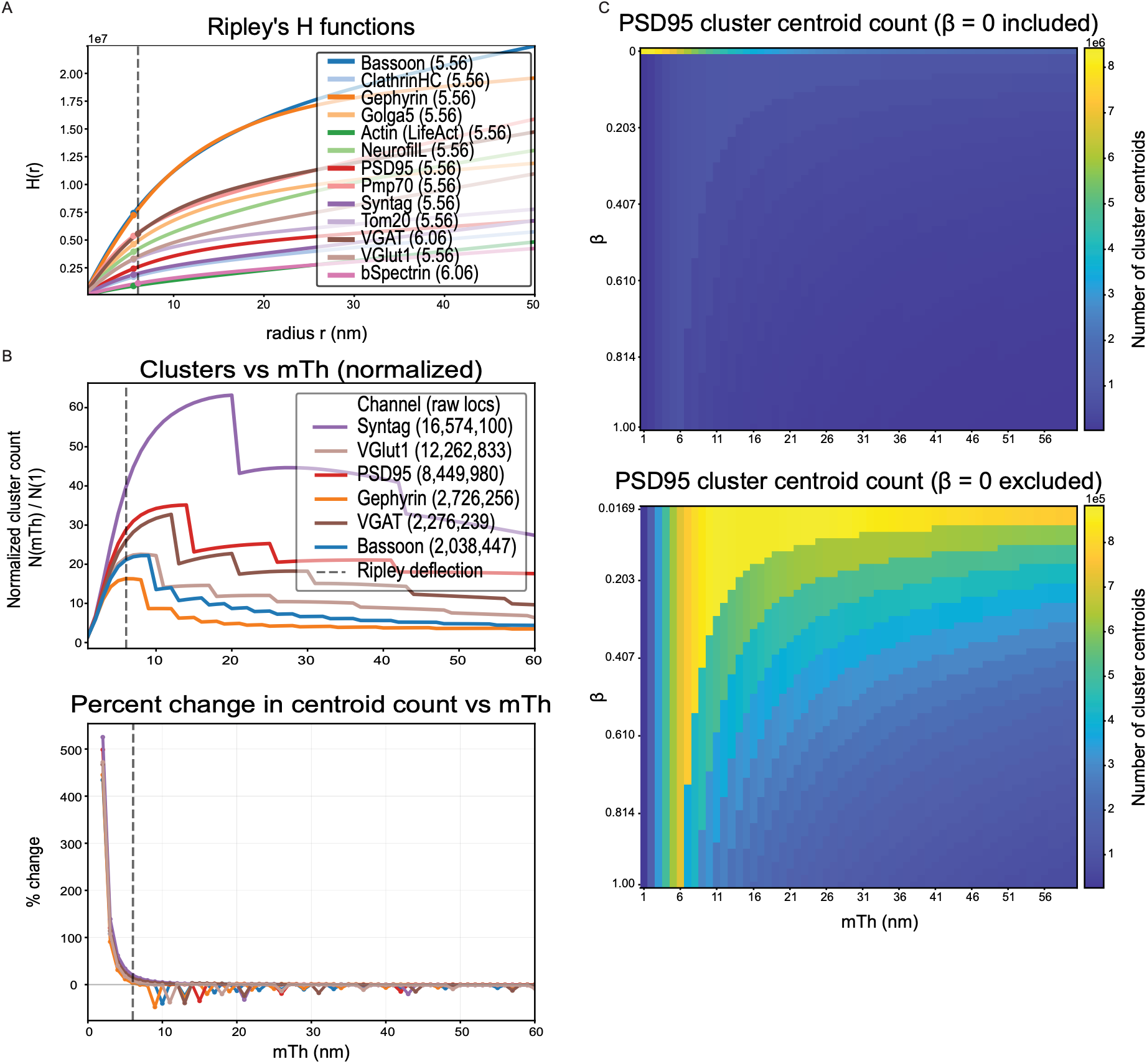
HDCluster parameter selection and sensitivity in experimental multiplexed DNA-PAINT data. (A) Ripley’s H analysis for the 13 DNA-PAINT channels^35^. The first small-scale deflection was detected at 5.56 nm for most channels and 6.06 nm for VGAT and βII-Spectrin. The dashed vertical line indicates the selected merging threshold, *mT*ℎ = 6 nm. (B) Effect of *mT*ℎ on cluster-centroid recovery for the six synaptic-marker channels across *mT*ℎ values from 1 to 60 nm. Top, cluster-centroid counts normalized to the count at *mT*ℎ = 1 nm, N(*mT*ℎ)/N(1); numbers in parentheses indicate the number of raw localizations in each channel. Bottom, percent change in centroid count relative to the immediately preceding *mT*ℎ value. The dashed vertical line indicates *mT*ℎ = 6 nm. (C) PSD95 cluster-centroid counts across combinations of *mT*ℎ from 1 to 60 nm and the optional denoising parameter β from 0 to 1. Top, analysis including β = 0; bottom, the same analysis with β = 0 excluded to better present differences among denoising-enabled conditions. Corresponding analyses for all six synaptic-marker channels are shown in Supp. Figure 6.

Using *mT*ℎ = 6 nm and the default denoising β = 0.2, we identified excitatory and inhibitory structures based on the alignment of VGlut1/Bassoon/PSD95 and VGAT/Bassoon/Gephyrin, respectively^35^, within an 800-nm-diameter sphere^40^. Detected excitatory and inhibitory structures within a representative ROI are shown in Figure 5A. The maximum number of both excitatory and inhibitory structures was detected around *mT*ℎ = 6 nm, and analysis of detected synapses showed that the number of centroids detected for each of the three marker channels, for both excitatory and inhibitory synapses, reached or approached maximum at *mT*ℎ = 6 nm (Figure 5B). Representative views of excitatory and inhibitory synapses further highlight preservation of the stratified three-marker organization at *mT*ℎ = 6 nm compared with lower and higher *mT*ℎ values (Figure 5C). The resulting Ripley-guided HDCluster processing and structure-recovery workflow is summarized in Figure 5D and compared with a published approach used to identify excitatory, inhibitory, and mixed synapses from similar DNA-PAINT data^40^. Importantly, in contrast to the application of DBSCAN to identify clusters within manually identified ROIs containing multiple excitatory or inhibitory synaptic markers^40^, the HDCluster pipeline automates the process, applying HDCluster across the full FOV for each marker channel and identifying synapses based on spatial organization of the resulting cluster centroids. This removes the need for manual ROI selection prior to clustering and allows cluster centroids generated across each full marker channel to be used directly for automated synapse detection.

**Figure 5.**
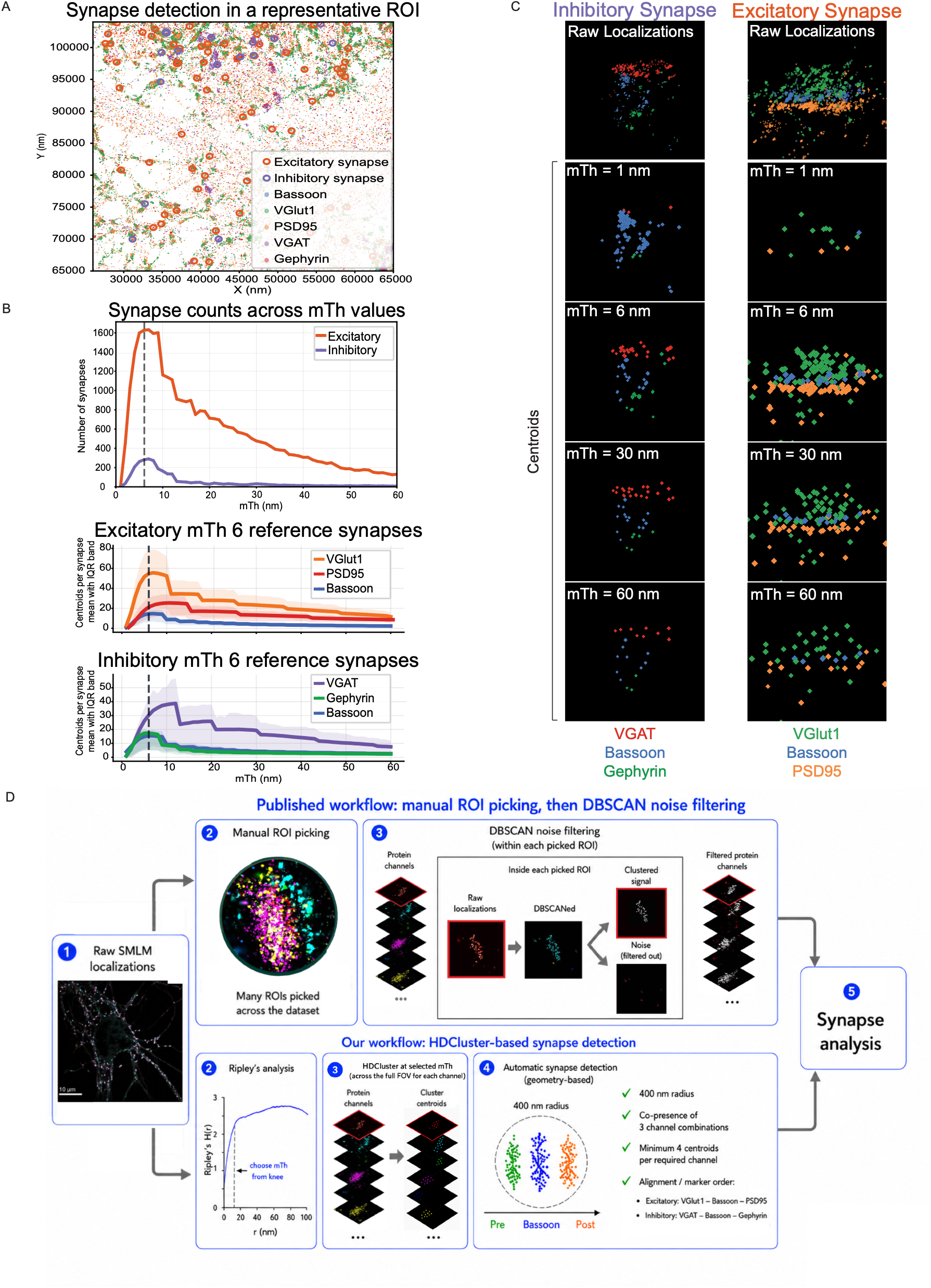
Effect of HDCluster *mT*ℎ on recovery of synaptic marker organization in experimental DNA-PAINT data. (A) Representative neuronal ROI showing synaptic-marker localizations and detected excitatory and inhibitory synapses at *mT*ℎ = 6 nm. Excitatory and inhibitory synapses were identified using VGlut1/Bassoon/PSD95 and VGAT/Bassoon/Gephyrin, respectively (see Methods for details)^35^. (B) Effect of *mT*ℎ on synapse detection and centroid recovery across *mT*ℎ values from 1 to 60 nm. Top, numbers of excitatory and inhibitory synapses detected after independently applying the same detection criteria at each *mT*ℎ. Middle and bottom, synapses detected at *mT*ℎ = 6 nm were retained as fixed reference populations, and the numbers of VGlut1, Bassoon and PSD95 centroids per excitatory synapse (middle) or VGAT, Bassoon and Gephyrin centroids per inhibitory synapse (bottom) were quantified across the *mT*ℎ sweep. Lines show the mean and shaded regions show the interquartile range. Dashed vertical lines indicate *mT*ℎ = 6 nm. (C) Representative inhibitory and excitatory synapses detected at *mT*ℎ = 6 nm. Raw localizations are shown together with HDCluster centroids obtained from the same spatial regions at *mT*ℎ = 1, 6, 30 and 60 nm. (D) Comparison of the previously published SUM-PAINT analysis workflow, involving manual synapse selection followed by DBSCAN-based noise filtering within each selected ROI^40,41^, with the Ripley-guided HDCluster workflow used here, in which the selected *mT*ℎ is applied across the full FOV for each marker channel to generate cluster centroids, followed by automated geometry-based synapse detection.

## CONCLUSION

HDCluster is a simple, scalable, one-parameter clustering method that is very fast and robust to noise. By combining density-based clustering with graph-based techniques, HDCluster provides robust and accurate cluster identification in SMLM datasets, even with very large datasets of hundreds of thousands or millions of localizations, which allows for precise emitter reconstruction. Application of HDCluster to the published multiplexed neuronal DNA-PAINT dataset further demonstrates its utility for complex experimental SMLM data^35^. The identification of a common Ripley-derived *mT*ℎ near 6 nm across channels, together with limited sensitivity to β around this scale and preservation of higher-order marker organization, supports the robustness of Ripley-guided HDCluster processing for experimental DNA-PAINT data.

## METHODS

### 1. HDCluster Algorithm

HDCluster is an unsupervised algorithm for spatial data clustering. It is an iterative similarity graph-based clustering method that discovers clusters and estimates their centroids, which are referred to as consensus localizations that approximate the true emitter positions. Figure 1A illustrates the algorithm based on 2D data points and details the underlying methodology (See Supplementary Videos for an animation of the HDCluster algorithm workflow). However, the algorithm works for 3D spatial data as well. The one parameter that the user needs to choose to run the algorithm is the “merging threshold (*mT*ℎ)”. More formally, HDCluster pseudocode is provided in Algorithm 1. The algorithm begins by constructing a similarity graph based on the spatial proximity parameter *mT*ℎ and calculating the node degree. Aside from the proximity (based on *mT*ℎ) there are no other restrictions imposed on the similarity graph construction. Nodes are sorted based on the degree in descending order, so the algorithm iteratively merges each high-degree node with its connected neighbours, computing their mean spatial coordinates. The algorithm keeps iterating until all nodes are visited and produces a set of mean nodes that represent preliminary cluster centers. In the next stage, HDCluster reconstructs a new graph using the mean nodes and repeats the merging process across multiple iterations until the convergence condition is reached, where no edges remain between mean nodes, which is an indication that clusters are fully separated. Each top-level mean node is then assigned a unique cluster ID, and labels are backtracked from these centroids to the original nodes such that every localization/point belongs to one cluster only. The resulting top-level mean nodes represent the emitters of the extracted nanocluster/cluster. Additionally, HDCluster can incorporate a denoising step to filter out low-degree nodes as described in Algorithm 2, effectively removing noisy/background localizations (see Methods Section 4 for a more extensive discussion of filtering options).

The iterative merging nature in HDCluster is deterministic and based on a fixed set of rules where, at each iteration, clusters are merged according to well-defined similarity and neighbourhood conditions, and the process terminates when no further merges satisfy these criteria. The algorithm guarantees convergence in a finite number of steps because the number of clusters strictly decreases after each valid merge. The procedure terminates when no additional pairs satisfy the merging conditions, producing the final partition.

The robustness of HDCluster was evaluated by introducing an isolated outlier between two Gaussian clusters (σ = 4 nm). While the perturbation narrowed the optimal merging threshold range from 7–17 nm to 10–15 nm due to intermediate network alterations, the algorithm successfully converged to the correct two-cluster assignment, demonstrating high stability against minor data variations. Step-by-step animations visualizing these iterations before and after the perturbation are provided as Supplementary Videos.

The *mT*ℎ parameter selection is based on the physical characteristics of the underlying clusters, and one of the heuristics to set this parameter is based on Ripley’s H function (see Methods Section 3 for a more extensive discussion of *mT*ℎ). However, it can be selected based on the application and the spread of the clusters, as illustrated in Supp. Figure 5 and Equation (2). β is an optional parameter to control the noise removal with a default setting that can be changed if needed (see Methods Section 4 for a more extensive discussion of β). HDCluster is available as a GUI-based software and MATLAB and Python based code for batch processing at https://github.com/NanoscopyAI/HDCluster.

### 2. Computational Complexity Comparison of HDCluster and DBSCAN

The computational complexity of the HDCluster can be derived from the pseudocode provided in Algorithm 1, where at each hierarchical merging level, HDCluster uses KD-tree range search to construct the radius-based similarity graph. For a dataset with *n* points, the KD-tree construction requires *O*(*n* log *n*) time, and within *mT*ℎ it performs radius queries for all points that require *O*(*n* log *n* + *nk*), where *k* indicates the average node degree in the neighbourhood graph (i.e., the average number of neighbours per point). Node degrees are computed in *O*(*n*) time, and nodes are sorted by degree in *O*(*n* log *n*). The subsequent merging processes each node and its local neighbourhood once, which requires *O*(*nk*) time. Therefore, the total time complexity per hierarchy level is *O*(*n* log *n* + *nk*).

As HDCluster iteratively merges points across *L* hierarchy levels, the overall complexity is 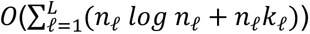, where *n*ℓ is the number of remaining representatives at level ℓ. Practically, for 3D point clouds with bounded local density (*k* ≪ *n*) and a small number of hierarchy levels, the runtime approaches *O*(*Ln* log *n*). In the worst case, if the neighbourhood size grows proportionally with the dataset (*k* = *O*(*n*)), the complexity degrades to *O*(*Ln*^2^), even though this scenario is uncommon for spatial point clouds.

On the other hand, DBSCAN revisits neighbourhoods during cluster expansion, which requires *O*(*n* log *n* + *nk* + *m*) time, where *m* = number of edge traversals. Since *m* ≈ *nk*, then the computational complexity of the DBSCAN requires *O*(*n* log *n* + *nk*) time. The worst case scenario for DBSCAN requires *O*(*n*^2^) time if all points are neighbours.

Although HDCluster and DBSCAN show similar theoretical complexity when implemented with KD-tree–based neighbourhood queries, their empirical runtimes differ substantially due to differences in how neighbourhoods are processed and how the dataset size evolves during clustering. Both methods require radius-based neighbour searches with expected complexity *O*(*n* log *n* + *nk*), where *n* is the number of points and *k* is the average neighbourhood size. However, DBSCAN performs recursive density expansion, repeatedly visiting neighbourhoods during cluster growth and maintaining additional bookkeeping for core, border, and visited points. These operations introduce significant overhead and limit opportunities for reducing the working set size. HDCluster performs a single degree-guided merging pass per hierarchy level, such that each point contributes only through its local neighbourhood. This arrangement reduces redundant neighbourhood traversals and enables efficient clustering of large-scale point clouds when combined with KD-tree–based range search (See Supp. Table 4).

### 3. HDCluster Merging threshold (*mT*ℎ) and Evaluation Measures

Merging threshold (*mT*ℎ) in HDCluster is a spatial proximity parameter that governs the similarity graph construction and guides the iterative merging of nodes toward the convergence of cluster means of a consensus localization. The selection of *mT*ℎ depends on the intrinsic properties of the dataset and the physical characteristics (i.e., size or dispersion) of the clusters. In experimental datasets, such as the SMLM data where ground truth is unavailable, *mT*ℎ can be estimated using spatial statistical analyses such as Ripley’s H function, where the clustering scale is identified from the peak clustering radius (i.e., the radius corresponding to the peak deviation from complete spatial randomness). Moreover, data-driven heuristics may be leveraged by examining the distribution of local connectivity and selecting a value that maintains cluster cohesion and simultaneously avoids the merging of distinct neighbouring clusters (Supp. Figure 5A). Because *mT*ℎ defines the maximum separation between highest-degree regions (i.e., density maxima) that may belong to the same structure, values near cluster scale generally provide robust performance. Additionally, our analysis (Figure 2C) shows that HDCluster maintains stable clustering performance over a wide range of *mT*ℎ values around the clustering scale, indicating that HDCluster is not highly sensitive to small changes in the *mT*ℎ parameter (See Supplementary Videos for an animation of the HDCluster and its sensitivity to parameter adjustments).

To demonstrate how we can set the *mT*ℎ parameter, we simulated various scenarios of neighbouring emitters (Supp. Figure 5A) with varying separation distance, defined as the space between the centers of the clusters. σ is the standard deviation of the Gaussian that quantifies the spread of the localizations/points around each cluster center. We derived the overlapping distance as a function of σ and separation distance to quantify the extent of overlap between the clusters, with negative values reflecting the separation between the boundaries of non-overlapping clusters (Equation (1)). The parameter *mT*ℎ can be set according to Equation (2). We also define three measures of error: *counting error (error_C), localization error (error_L1),* and *localization error (error_L2)* (Equations (3–5)).

To assess the applicability of the HDCluster in reconstructing the emitters, we simulate various scenarios that may be similar to real-world situations, where the molecules can be sparse or very dense. We simulated two Gaussian emitters and change two parameters, (1) the separation distance (i.e., the distance between the centers/means of the two Gaussians or the emitters, which we use to simulate the density of the molecule in larger grids) and (2) the standard deviation of the Gaussians σ, that simulates the localization precision, i.e., the uncertainty in detecting the position of that emitter across repeated localizations, as shown in Supp. Figure 5A. The overlapping distance between the two clusters is defined as the distance between the borders of the clusters. The *overlap* between the localizations of the emitters is calculated according to Equation (1). The overlapping distance is a function of σ and *separation* distance. In the simulation, we can change the *separation* distance and σ to compute the amount of *overlap* between the neighbouring clusters. The *overlap* is negative when the localizations of the neighbouring emitters are not overlapping, and the *overlap* is positive when the localizations of the neighbouring emitters are overlapping. The *overlap* equals zero when the borders of the two clusters are touching each other (Supp. Figure 5A).

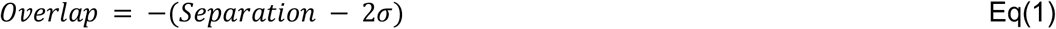

Where σ is the standard deviation of the Gaussian-generated data.

The HDCluster parameter, *merging threshold (mT*ℎ), can be selected to reconstruct the emitters’ locations based on Equation (2).

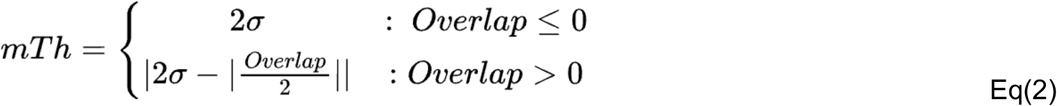

In Equation (2), when the neighbouring clusters are well separated or their borders touch each other (i.e., *Overlap* ≤ 0), the parameter *mT*ℎ = 2*σ* ensures appropriate merging within each cluster. However, as clusters become closer and their borders overlap (i.e., *Overlap* > 0), *mT*ℎ should be reduced to prevent unintended merging of adjacent overlapping clusters (i.e., *mT*ℎ can be safely adjusted to 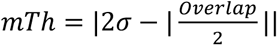. This approach aligns with adaptive thresholding strategies used in hierarchical and graph-based clustering methods, where thresholds decrease as clusters become denser or closer.

More formally, we simulated 3D grids of size 3 × 3 × 3 and we changed the *separation* distance and the standard deviation of the Gaussians σ. Moreover, we also changed the density of the localizations around everyone of the simulated emitters as shown in Supp. Figure 5B. HDCluster is used to reconstruct the emitters. The parameter *mT*ℎ is selected according to Equation (2). To assess the quality of emitter location reconstruction, we introduced three measures, as we have the ground truth location for the emitters. We quantitatively calculated three types of errors: *error_C* Equation (3), *error_L1* Equation (4), and *error_L2* Equation (5).

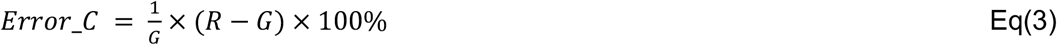

where *G* is the number of ground truth (GT) emitters and *R* is the reconstructed emitters or binding sites. In the *error_C*, positive values mean overcounting, negative values mean undercounting, and zero means no counting error.

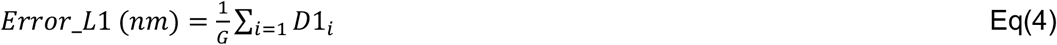

where *D*1*_i_* is the distance between the GT emitter *i* and the location of the nearest reconstructed emitter. *Error*_*L*1 is normalized by the number of ground truth emitters *G*.

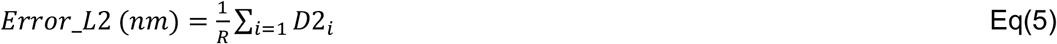

where *D*2*_i_* is the distance between the reconstructed emitter *i* and the location of the nearest GT emitter. *Error*_*L*2 is normalized by the number of reconstructed emitters *R*.

We generated a larger 3D 3 × 3 × 3 grid of simulated emitters and localizations, where the localizations’ membership and the emitters’ location are considered as a ground truth (GT) for evaluation via the error measures (Supp. Figure 5B). We generated 20 grids for each pair of σ and separation distance and computed the three types of counting (i.e., *error_C*) and localization errors (i.e., *error_L1* and *error_L2*). We started with a low density of 10 localizations per emitter and increased the density to 35 localizations per emitter. We notice that the counting error (*error_C*) is improved (Supp. Figure 5B, left vs right), indicating emitter reconstruction is much better for a higher density of localizations. As σ decreases (i.e., more precise localizations), the 3 types of error become smaller (Supp. Figure 5B). For example, when using *σ* = 2, for any separation distance ≥10 nm, and for a number of localizations 35 per emitter, the *error*_*C* is zero, and both the *error*_*L*1 and *error*_*L*2 are almost zeros (i.e., the location of the reconstructed emitters is almost identical to the ground truth location). The error graphs show that the reconstruction errors remain relatively low, even under challenging conditions where nearby emitters exhibit substantial overlap. In the worst cases, *error*_*L*1 and *error*_*L*2 stay below ∼30 nm and 14 nm, respectively. *Error*_*L*1 and *error*_*L*2 fluctuate due to overcounting or undercounting, specifically in challenging cases of highly overlapping or low-density nanoclusters.

### 4. HDCluster Denoising Parameter - Spatial Filtering Option

HDCluster incorporates a denoising option to filter out noisy localizations (background, outlier points, etc.). The denoising capability/function in HDCluster can be enabled or disabled by the user. If enabled, the user can set a β value or rely on its default value. The embedded filtering works by calculating the degree for all the nodes, then labeling all the nodes with a degree below a *low degree* with a 0 label (i.e., noisy localizations), as explained in Algorithm 2. The *low_degree* value is calculated as shown in Equation (6).

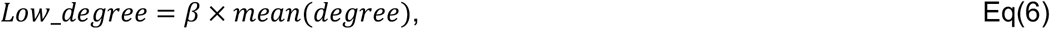

where *mean*(*degree*) is the average node degree that is calculated when the graph is constructed with the *mT*ℎ parameter, β which is a scalar that controls the denoising process. As a rule-of-thumb, the default value β = 0.2 is selected to label the non-clustered localizations as noisy. However, the user can select other β values that might control the noise removal according to the data, where larger β values employ harsh filtering. β is a scaling factor that controls the minimum node degree relative to the average connectivity of the graph, which determines how aggressively HDCluster removes low-connectivity nodes (potential noise). Setting β = 0.2 ensures that only nodes whose connectivity is <20% of the mean graph degree are labeled as noise and generalizes well across different datasets without prior tuning. Setting β balances between retaining true cluster points at the periphery and filtering isolated/background points that do not contribute meaningfully to the cluster structure. We provided the user with the ability to change β. Higher values (i.e., >0.2) can lead to harsher filtering, which can fragment clusters and remove localizations within low-density regions.

### 5. Generating Data with Various Shapes and Densities with Additive Noise

To test the robustness of the clustering methods to noise, we generated a dataset with *K* = 80 Gaussian clusters that varied in density and were randomly positioned in 3D space. Each cluster has a minimum of 10 points and a maximum of 100 (for various-density clusters), and of various shapes, varying the covariance matrix diagonal of the Gaussians as given in Equation (7).

Let each cluster *i* ∈ {1,2, … , *K*} be defined by:

- Mean *μ_i_* = [*μ_ix_*, *μ_iy_*, *μ_iz_*]*^T^* ∈ *R*^3^
- Covariance matrix (diagonal): 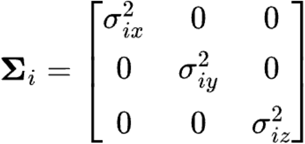

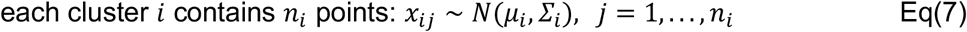

Sampling procedure:

(i) For each cluster *i* ∈ {1,2, … , *K*}, randomly sample the mean:

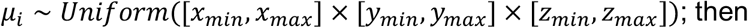
(ii) randomly sample diagonal variances:

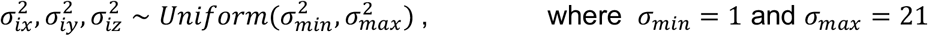

from the diagonal covariance matrix Z*_i_* as shown above. Sample *n_i_* points from the 3D Gaussian distribution *x_i_*_j_ ∼ *N*(*μ_i_*, Z*_i_*). Notice that the off-diagonal elements are all zero, so the clusters are axis-aligned ellipsoids. The variance values control the spread along each axis independently to get different shapes. We simulated the irregular cluster sizes by drawing *n_i_* randomly (i.e., uniform distribution) to vary the clusters’ densities. We then varied the noise by controlling the noise level, where the noise level can be selected for every experiment and drawn from a uniform distribution. HDCluster can effectively handle non-isotropic (ellipsoidal) cluster shapes, as demonstrated using Gaussian clusters with varying covariance matrices; however, it is not specifically designed for highly fibrous or filamentous structures, where cluster morphology deviates significantly from compact density-based regions.

### 6. Processing DNA-PAINT data with HDCluster

Published 13-plex neuronal DNA-PAINT data from Unterauer et al.^35^ were analyzed using HDCluster. For each of the 13 channels, Ripley’s H analysis was used to estimate the characteristic spatial scale of local multiple blinking localization networks. Ripley’s H curves were analyzed using a custom script to identify the initial small-scale deflection point, and these values were used to guide selection of the merging threshold parameter, *mT*ℎ. Because the detected deflections occurred near 6 nm across channels, *mT*ℎ = 6 nm was selected for the primary analysis. Unless otherwise stated, the default denoising parameter β = 0.2 was used. To assess sensitivity to *mT*ℎ, the data were additionally processed across *mT*ℎ values from 1 to 60 nm.

Cluster-centroid counts were quantified across the *mT*ℎ sweep for the six synaptic-marker channels (Bassoon, VGlut1, PSD95, VGAT, Gephyrin and Synaptotagmin1). Counts at each *mT*ℎ were normalized to the number obtained at *mT*ℎ = 1 nm, N(*mT*ℎ)/N(1). The percent change in centroid number between consecutive *mT*ℎ values was calculated relative to the immediately preceding *mT*ℎ as 100 × [*N*(*mT*ℎ*_i_*) − *N*(*mT*ℎ_(*i*−1)_)]/*N*(*mT*ℎ_(*i*−1)_). The effect of the optional denoising parameter β was assessed by varying β from 0 to 1 across *mT*ℎ values from 1 to 60 nm. Cluster-centroid counts for each *mT*ℎ–β combination were visualized as heatmaps both including and excluding β = 0.

Putative excitatory and inhibitory synapses were identified from HDCluster centroids using VGlut1/Bassoon/PSD95 and VGAT/Bassoon/Gephyrin, respectively^35^. Bassoon centroids were used as seed points, and neighbouring Bassoon, presynaptic (VGlut1, VGAT) and postsynaptic (PSD95, Gephyrin) centroids were collected within a 400-nm radius of each seed, corresponding to an 800-nm-diameter sphere^40^. Candidate synapses were required to contain at least four centroids from each required marker channel to reduce detections driven by sparse or isolated centroids. The pre-to-postsynaptic axis was defined from the center of the presynaptic-marker distribution to the center of the postsynaptic-marker distribution. Candidates were then filtered according to the expected spatial layering of the three marker classes along this axis. At least 70% of presynaptic-marker centroids were required to occupy the presynaptic side, at least 70% of Bassoon centroids the intermediate region, and at least 70% of postsynaptic-marker centroids the postsynaptic side. Bassoon centroids were additionally required to lie within 150 nm of the pre-to-postsynaptic axis. Accepted candidate centers were required to be at least 300 nm apart, and centroids assigned to an accepted candidate were not reused in subsequent candidates. The same detection criteria were consistently applied independently at each *mT*ℎ value to quantify excitatory and inhibitory synapse recovery across the *mT*ℎ sweep. Synapses detected at *mT*ℎ = 6 nm were retained as fixed reference ROIs, and the number of centroids from each of the three required marker channels was quantified within the same ROIs across *mT*ℎ values. Centroid numbers within the reference ROIs were plotted as the mean with interquartile range. Representative inhibitory and excitatory synapses detected at *mT*ℎ = 6 nm were selected for visualization of the raw localizations and HDCluster centroids from the same ROIs at *mT*ℎ = 1, 6, 30 and 60 nm.

## Supporting information

Various clustering methods applied to benchmarking shape datasets of a known number of clusters and points' memberships to which cluster.

Qualitative analysis comparison of the clustering methods based on the real SMLM dataset.

Applying HDCluster to reconstruct the binding sites from dSTORM nanoruler and DNA-PAINT origami nanostructures datasets.

Generation and performance evaluation of synthetic 3D Gaussian clusters for reconstruction error analysis.

## Acknowledgements

The authors gratefully acknowledge Dr. Johannes Stein (Harvard Medical School; currently, Max Planck Institute for Molecular Genetics) for his valuable discussions and for providing the DNA-PAINT dataset used in this study. We thank Dr. Antonios Lioutas for his valuable discussions and for providing infrastructure and experimental support for testing HDCluster. We also thank Dr. Ricardo Nunes Bastos (ONI) for kindly providing the Gatta-PAINT 80 nm nanoruler dataset acquired using the ONI Nanoimager and Eduard Unterauer for providing the 13-plex neuronal DNA-PAINT dataset. We also thank Dr. Ben Cardoen for developing the Ripley’s H knee-detection script and the Digital Research Alliance of Canada for computational resources. Funding for this project was provided by the Canadian Institutes of Health Research (CIHR: PJT-175112) and the Natural Sciences and Engineering Research Council of Canada (NSERC: RGPIN-2019-05179, RGPIN-2020-06752). Antonios Lioutas and Johannes Stein were supported by awards to Dr. C.-ting Wu from the National Institutes of Health (NHGRI; RM1HG011016 and UM1HG011593) and the Genome Imaging Fund of Harvard Medical School. I.M.K. gratefully acknowledges the support of Birzeit University and the Fulbright Visiting Scholar Program for funding this research, and Dr. C.-ting Wu and the Genome Imaging Fund of Harvard Medical School for hosting and supporting the completion of this work.

## Author Contributions

**I.M.K.** Conceptualization, methodology, software (core code and GUI), data gathering, data generation, data curation, formal analysis, investigation, visualization, writing - original draft.

**S.B.B.** Investigation (testing HDCluster on real DNA-PAINT datasets and 13-plex neuronal labeling experiment), formal analysis (assessing denoising parameter β), data visualization (Figure 4, Figure 5, and Supplementary Figure 6).

**S.J.E.** Software (Python translation), formal analysis (execution time benchmarking), data visualization and curation (Supplementary Figure 2 and Supplementary Table 4, including the LifeAct actin dataset).

**G.H. and I.R.N.** Project supervision, funding acquisition, methodology (G.H. contributed to the design and coding of the original iterative merging algorithm and code testing), writing - review & editing.

All authors reviewed and approved the final version of the manuscript.

## Data Availability

All datasets analysed in this study are either publicly available or can be regenerated using the scripts provided in our repository, as detailed below.

1. **Shape benchmark datasets.** The Aggregation, Flame, R15 and D31 shape sets^19^ used in Figure 1C and Supplementary Figure 1 are publicly available from the clustering benchmark collection of the University of Eastern Finland at https://cs.joensuu.fi/sipu/datasets/
2. **Simulated SMLM benchmark datasets.** The simulated SMLM point-cloud benchmark of Nieves et al.^26^, comprising 10 scenarios of 50 point-cloud files each, in ground-truth and multiple-blinking variants, is publicly available at https://github.com/DJ-Nieves/ARI-and-IoU-cluster-analysis-evaluation. The present work analysed Scenarios 2–10; Scenario 1 was excluded as a null-hypothesis control.
3. **13-plex neuronal DNA-PAINT dataset.** The multiplexed neuronal DNA-PAINT data of Unterauer et al.35 are deposited at Zenodo under https://doi.org/10.5281/zenodo.17023207. Both the six synaptic-marker channels analysed in Figures 4 and 5 (Bassoon, VGlut1, PSD95, VGAT, Gephyrin and Synaptotagmin1) and the LifeAct actin channel used for the large-scale runtime benchmark (65,910,759 localizations; Supplementary Table 4) are taken from this dataset.
4. **DNA origami nanostructures.** The linear and triangular DNA origami nanostructure data of Masullo et al.^38^ used in Supplementary Figure 4B are deposited at Zenodo under https://doi.org/10.5281/zenodo.15131642.
5. **DNA origami 3 × 4 grids.** The 3 × 4 DNA origami grid dataset with 20 nm site spacing used in Figure 3 was generated in Stein et al.^39^ and provided to us by the authors of that study.
6. **Gatta-PAINT 80 nm nanoruler.** The nanoruler dataset used in Supplementary Figure 4A was acquired on an ONI Nanoimager and provided by ONI (Oxford Nanoimaging Ltd). It is not deposited in a public repository; enquiries regarding access should be directed to ONI. The nanoruler sample itself is a commercial product of GATTAquant GmbH.
7. **HeLa caveolin-1 dataset.** The caveolin-1 dataset from HeLa cells (1,535,678 localizations) used in Figure 2E and Supplementary Figure 3 was previously published in Khater et al.^34^ and is available from the corresponding author upon reasonable request.
8. **Simulated Gaussian cluster data.** The 3D data simulated for this study — the 80-cluster datasets with additive uniform noise (Methods §5; Figure 2) and the 3 × 3 × 3 emitter grids used for reconstruction-error analysis (Methods §3; Supplementary Figure 5) — are not deposited as files but can be regenerated exactly using the simulation scripts at https://github.com/NanoscopyAI/HDCluster.

## Code Availability

HDCluster is available as GUI-based MATLAB software and as MATLAB and Python executable functions/scripts for batch processing at https://github.com/NanoscopyAI/HDCluster, together with the simulation and evaluation scripts used in this study.

## Algorithms

### Algorithm 1 HDCluster: High-Degree Graph Clustering

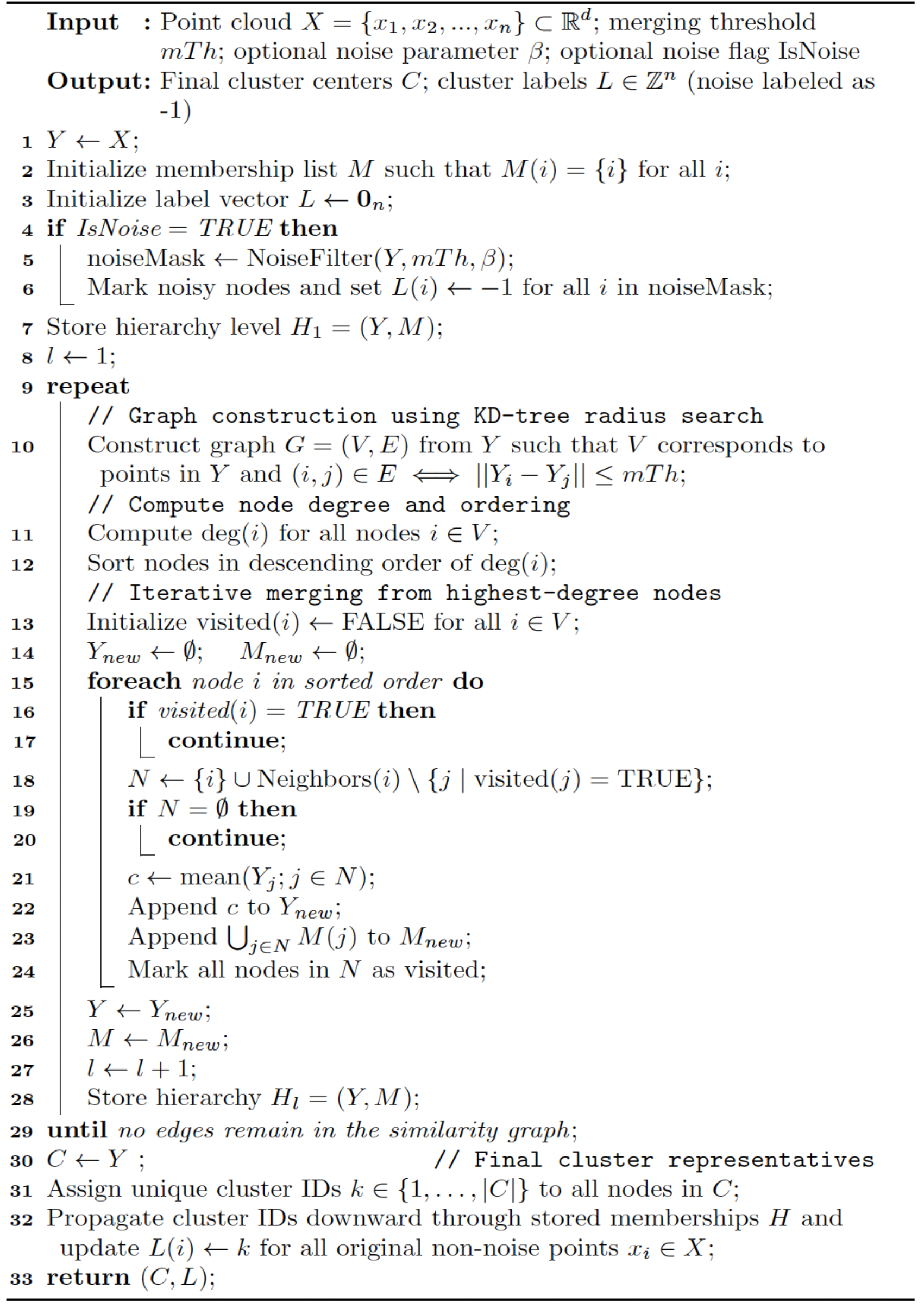

### Algorithm 2 NoiseFilter: Degree-Based Denoising

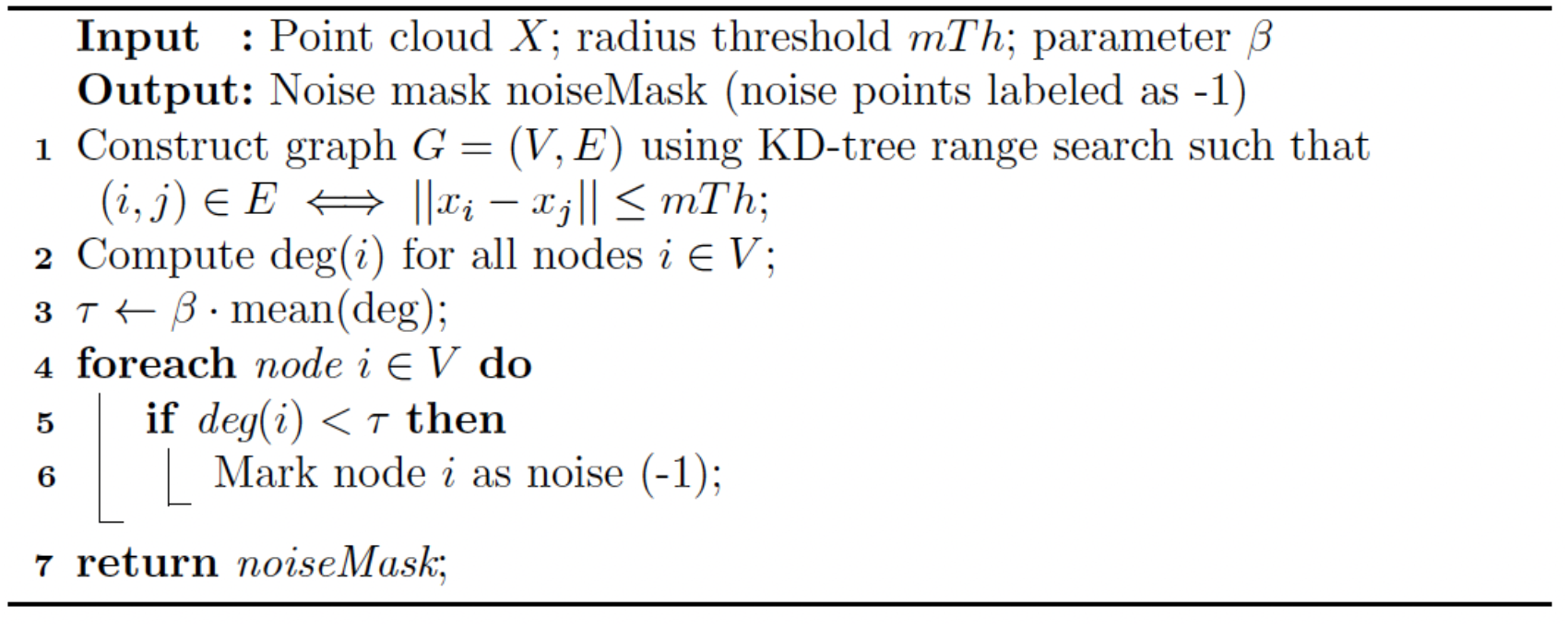

## Supplementary Information

**Supp. Table 1:**
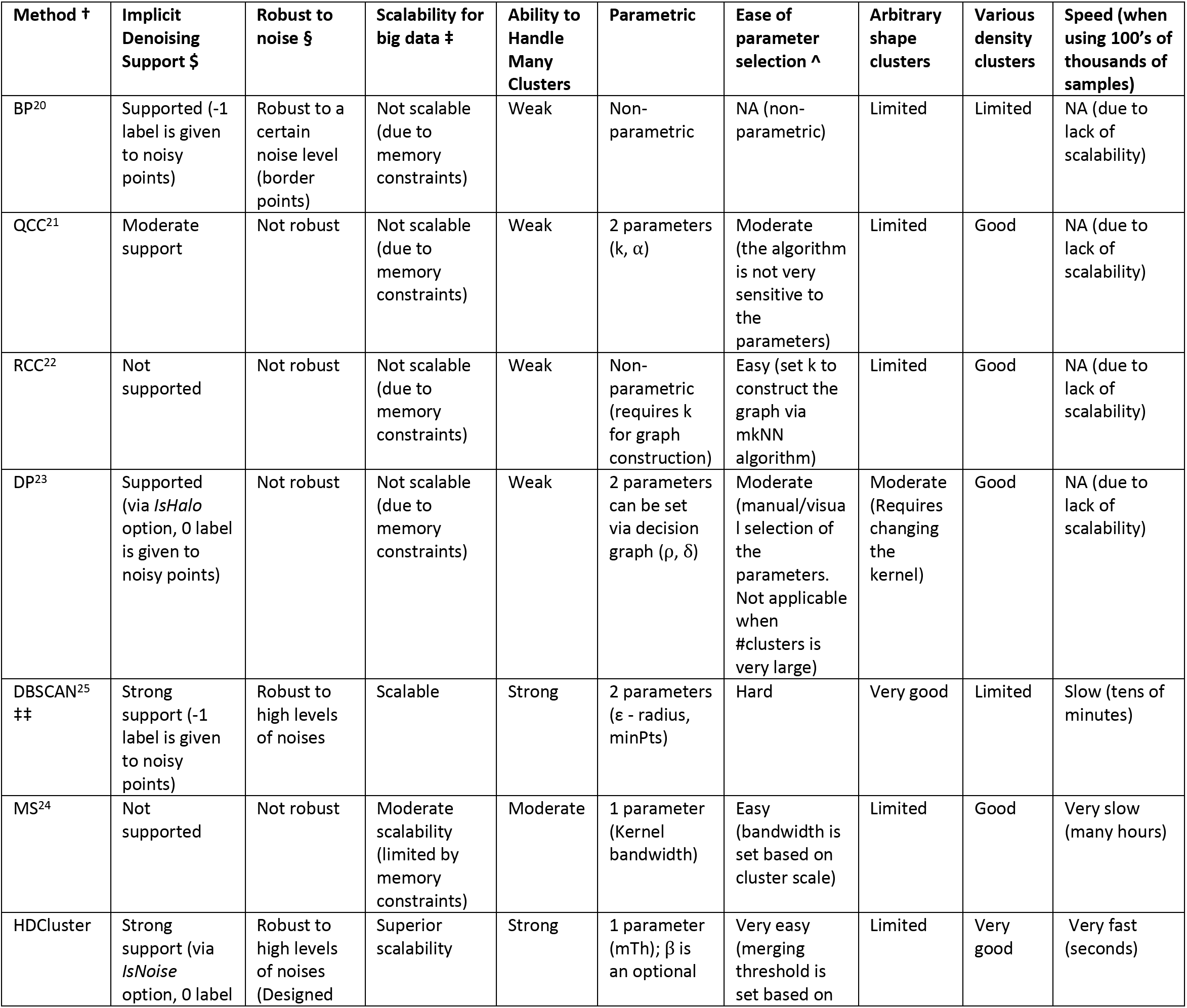
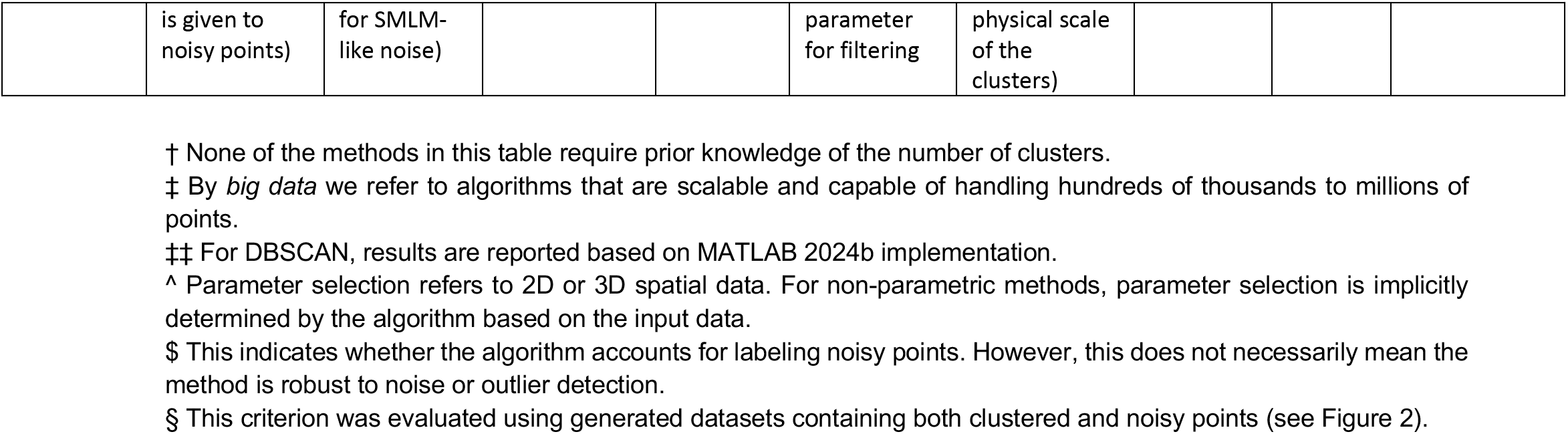
Comparison of state-of-the-art spatial clustering methods^20–25^ based on their key characteristics and performance criteria. The table lists state-of-the-art clustering methods with their characteristics, with a particular focus on their applicability to SMLM data. Based on our study, we enumerate each method’s robustness to noise, ability to explicitly label noisy localizations or points, scalability to large datasets, and effectiveness when dealing with a high number of clusters. We also assess the number of required parameters, the ease of parameter tuning, the ability to detect clusters of varying shapes and densities, and the computational efficiency when applied to large datasets containing hundreds of thousands of points.

**Supp. Table 2:** Summary of clustering evaluation metrics across various scenarios and methods. Results are presented as the mean ± standard deviation, calculated from 50 independently generated files for each scenario in SMLM benchmarking dataset^26^. DBSCAN adopts the parameters as reported in Nieves et al.^26^ that were tuned based on ARI. Look at Supp. Table 3 for the parameters used to generate this table. Where, SEM is the standard error of the mean (SEM = std / sqrt(N)), where N=9 scenarios.

|  | Method | Ground Truth (GT)<br>(No Added Blinking) |  | Multiple Blinking (MB) |  |
| --- | --- | --- | --- | --- | --- |
|  |  | ARI | AMI | ARI | AMI |
| Scenario 2 | HDCluster | **0.86 ± 0.02** | **0.85 ± 0.018** | **0.74 ± 0.059** | **0.82 ± 0.03** |
| | DBSCAN | 0.85 ± 0.023 | 0.84 ± 0.02 | 0.73 ± 0.061 | 0.82 ± 0.034 |
| Scenario 3 | HDCluster | **0.90 ± 0.017** | **0.83 ± 0.025** | **0.72 ± 0.057** | **0.71 ± 0.045** |
| | DBSCAN | 0.84 ± 0.017 | 0.74 ± 0.022 | 0.69 ± 0.05 | 0.69 ± 0.042 |
| Scenario 4 | HDCluster | **0.68 ± 0.064** | **0.62 ± 0.055** | 0.34 ± 0.099 | 0.41 ± 0.079 |
| | DBSCAN | 0.67 ± 0.057 | 0.60 ± 0.057 | 0.34 ± 0.103 | 0.41 ± 0.084 |
| Scenario 5 | HDCluster | 0.55 ± 0.027 | **0.72 ± 0.009** | 0.38 ± 0.036 | **0.65 ± 0.012** |
| | DBSCAN | **0.56 ± 0.02** | 0.71 ± 0.011 | **0.42 ± 0.034** | 0.64 ± 0.018 |
| Scenario 6 | HDCluster | 0.70 ± 0.02 | 0.74 ± 0.012 | 0.63 ± 0.036 | 0.70 ± 0.018 |
|  | DBSCAN | **0.73 ± 0.017** | **0.75 ± 0.016** | **0.65 ± 0.027** | **0.71 ± 0.018** |
| Scenario 7 | HDCluster | **0.55 ± 0.041** | **0.61 ± 0.027** | 0.43 ± 0.074 | 0.64 ± 0.032 |
| | DBSCAN | 0.53 ± 0.043 | 0.59 ± 0.036 | **0.45 ± 0.062** | 0.64 ± 0.032 |
| Scenario 8 | HDCluster | **0.82 ± 0.034** | **0.82 ± 0.03** | 0.64 ± 0.067 | **0.76 ± 0.044** |
| | DBSCAN | 0.80 ± 0.037 | 0.81 ± 0.031 | 0.64 ± 0.061 | 0.74 ± 0.062 |
| Scenario 9 | HDCluster | 0.57 ± 0.026 | **0.65 ± 0.014** | 0.54 ± 0.037 | **0.63 ± 0.02** |
| | DBSCAN | **0.58 ± 0.021** | 0.63 ± 0.02 | **0.55 ± 0.038** | 0.61 ± 0.028 |
| Scenario 10 | HDCluster | **0.85 ± 0.026** | 0.84 ± 0.025 | 0.71 ± 0.065 | 0.81 ± 0.034 |
| | DBSCAN | 0.84 ± 0.028 | 0.84 ± 0.024 | **0.72 ± 0.066** | **0.82 ± 0.034** |

| Metric | HDCluster<br>(Mean ± SEM) | DBSCAN<br>(Mean ± SEM) | Absolute diff.<br>(HDCluster – DBSCAN) | Relative Improvement (%)<br>$= \frac{HDCluster\ Mean - DBSCAN\ Mean}{DBSCAN\ Mean} \times 100$ |
| --- | --- | --- | --- | --- |
| GT ARI | 0.72 ± 0.05 | 0.71 ± 0.04 | 0.01 | 1.41 % |
| GT AMI | 0.74 ± 0.03 | 0.72 ± 0.03 | 0.02 | 2.78 % |
| MB ARI | 0.57 ± 0.05 | 0.577 ± 0.05 | – 0.007 | – 1.16 % |
| MB AMI | 0.68 ± 0.04 | 0.676 ± 0.04 | 0.01 | 0.83 % |

**Supp. Table 3:**
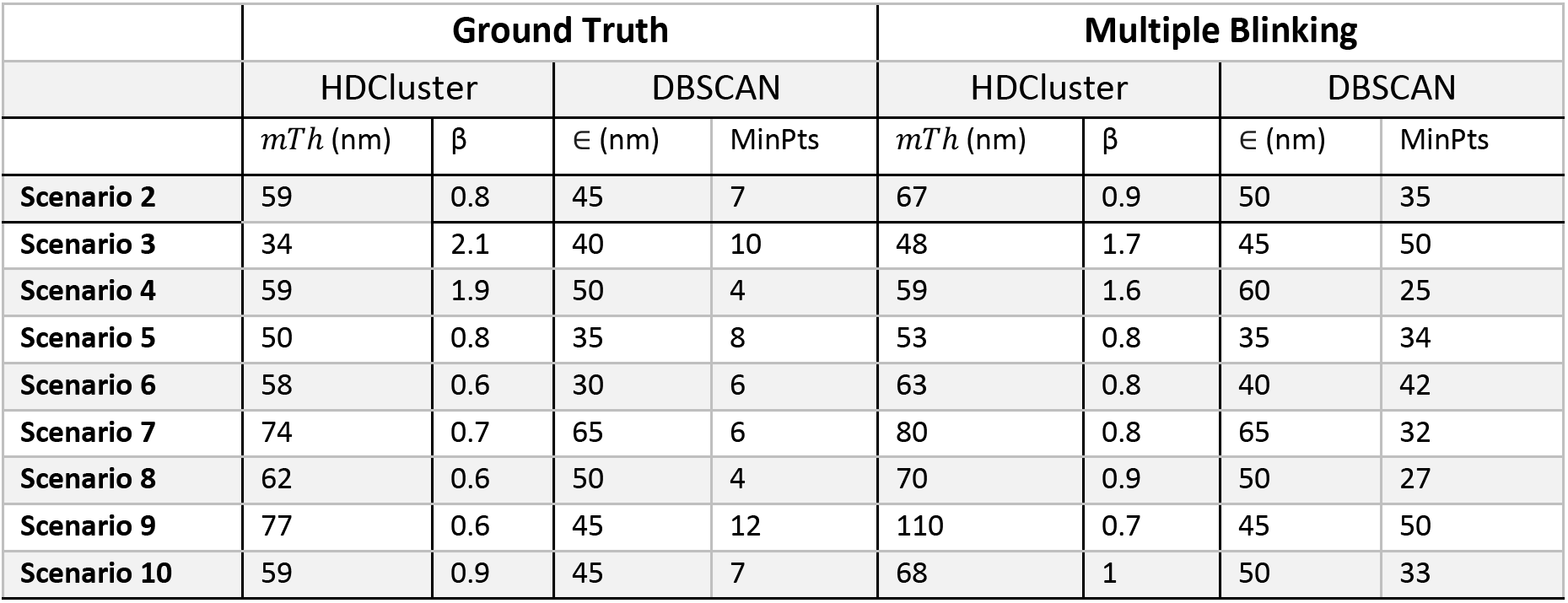
Methods’ Parameters. DBSCAN parameters adopted from Nieves et al.^26^. The HDCluster parameters are tuned based on the AMI measure.

**Supp. Table 4:**
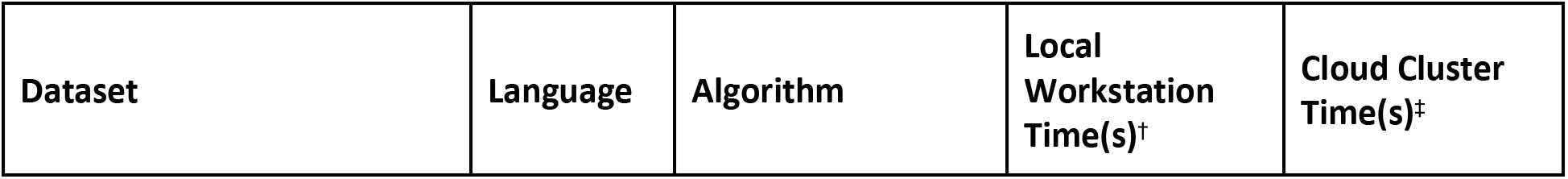
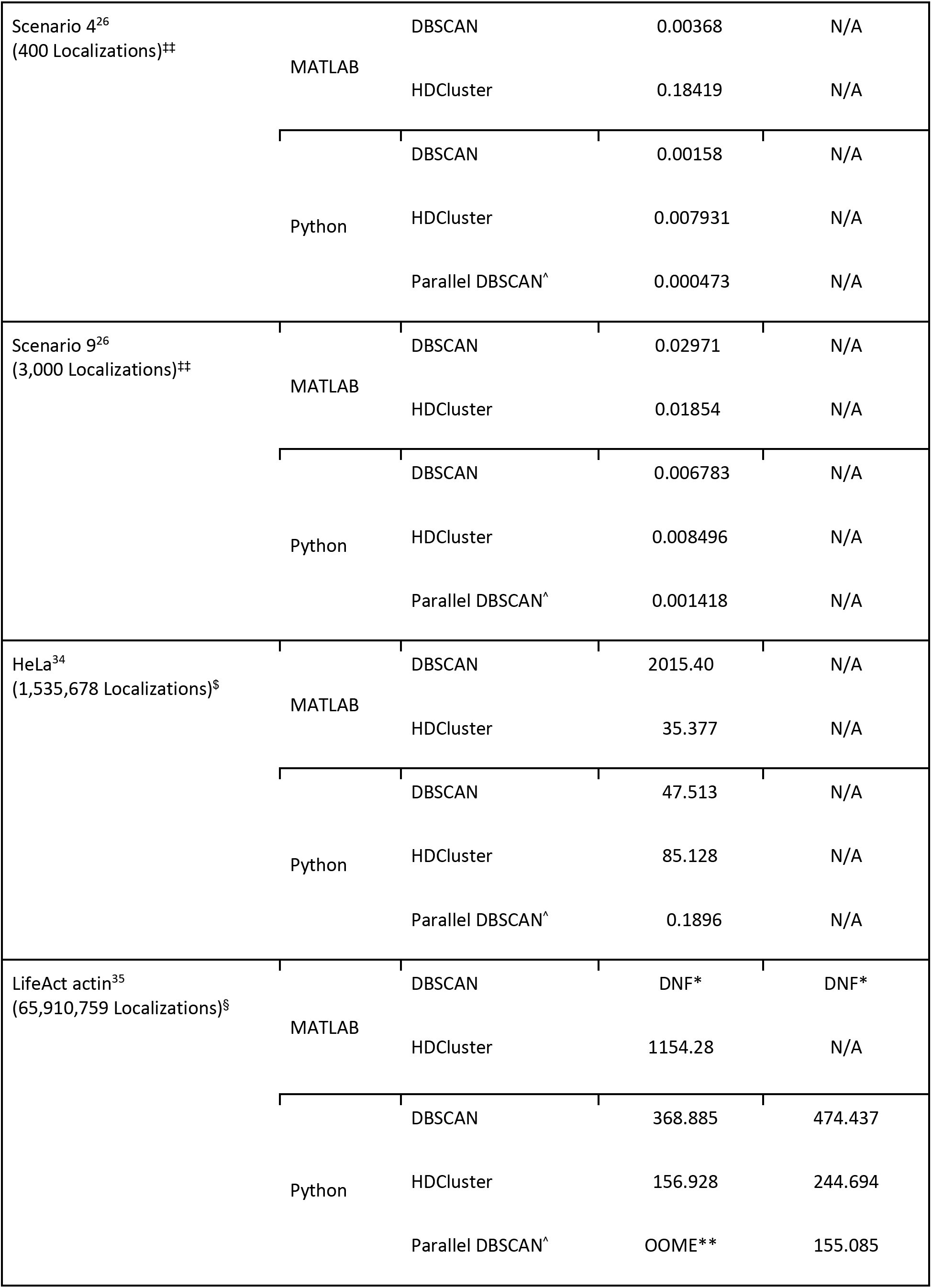
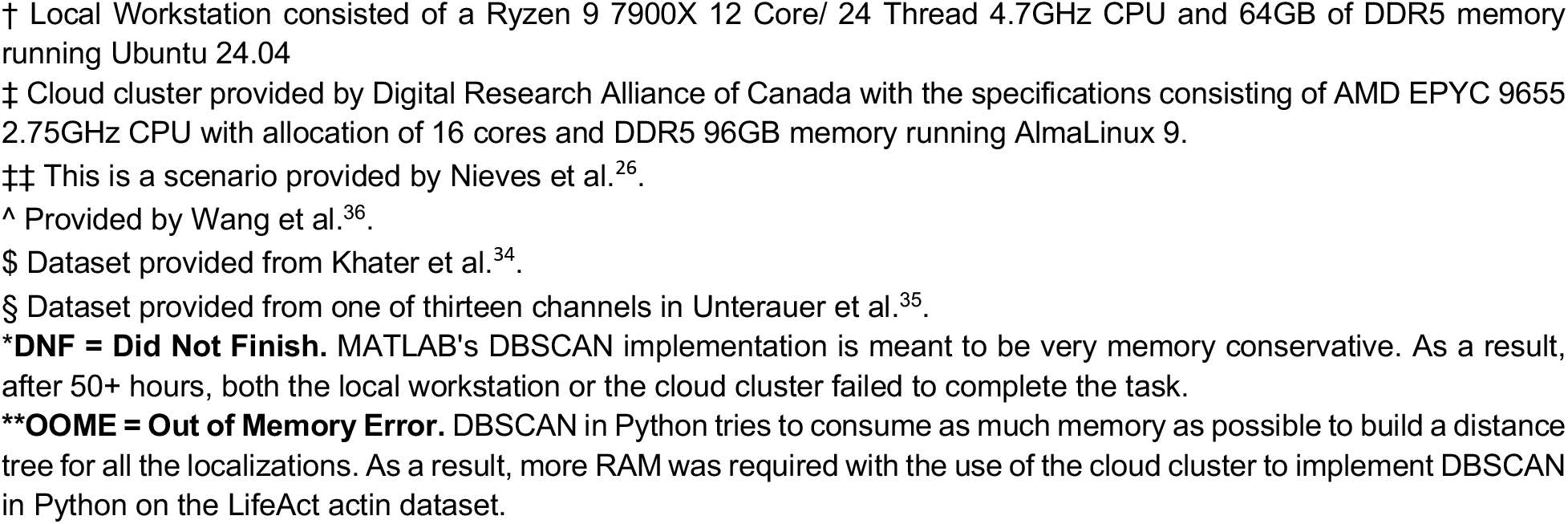
The compute times of HDCluster, and the most competitive clustering algorithm for SMLM data, DBSCAN. Compute times were acquired using the optimal parameters in Supp. Table 3. Scenarios 4 and 9 were chosen for comparison with Nieves et al.^26^ results. DBSCAN for Python displays high performance due to maturity in implementation, including the parallelization approach adopted by Wang et al^36^. As a result, the performance was high for a smaller sized dataset. For users working in MATLAB, HDCluster showed faster clustering times. When working with very large datasets such as LifeAct actin labeling by DNA-PAINT^35^, HDCluster is shown to be robust on the local workstation, where DBSCAN required a higher amount of memory to process its algorithm. This demand of memory requires a high memory workstation or to make use of cloud compute resources. Even with the help of the Digital Research Alliance of Canada cloud cluster, HDCluster was shown to perform faster than DBSCAN.

**Supplementary Video 1. Network construction and convergence of HDCluster on the baseline dataset.** This animation visualizes the step-by-step iterations of the HDCluster algorithm, including network construction, node degree calculations, local mean convergence, and backtracking-based cluster assignment, on a synthetic dataset consisting of two Gaussian clusters (σ = 4 nm, 15 localizations each) with the denoising option disabled. The clustering results are shown across a sequence of merging thresholds (*mT*ℎ = 5, 7, 10, 15, 17, and 20 nm), demonstrating accurate two-cluster retrieval within the stable range of 7 to 17 nm, while under-segmentation and over-segmentation occur at the extreme thresholds of 5 nm and 20 nm, respectively. https://youtu.be/lxGGBm4I67I?si

**Supplementary Video 2. Algorithmic robustness and convergence of HDCluster under minor data perturbation.** This animation illustrates the iterative behavior and clustering performance of HDCluster following the introduction of a single outlier point between the two baseline clusters at coordinates [−17, −35]. Using the same sequence of merging thresholds (*mT*ℎ = 5, 7, 10, 15, 17, and 20 nm), the video demonstrates how the perturbation alters the local network topology at intermediate thresholds (e.g., resulting in three clusters at *mT*ℎ = 7 nm). Despite a narrower optimal parameter window (*mT*ℎ = 10–15 nm), the algorithm successfully converges to the correct two-cluster assignment, highlighting its robustness against minor data variations. https://youtu.be/6l_F9WLsUVQ?si

### Supplementary Figures

**Supplementary Figure 1.**
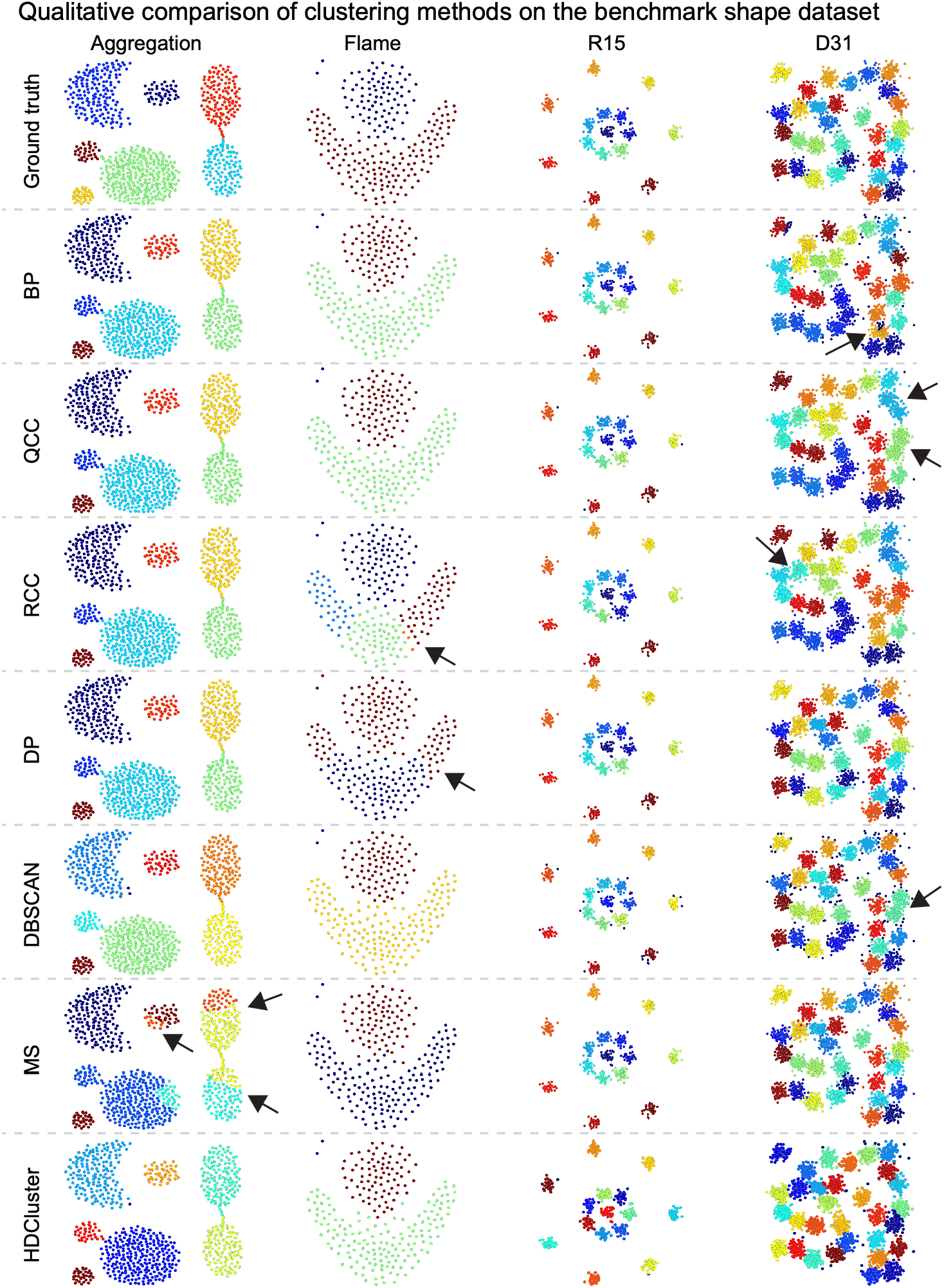
Various clustering methods applied to benchmarking shape datasets of a known number of clusters and points’ memberships to which cluster. The qualitative results comparison of the various clustering methods shows how the methods classify the points in comparison to the provided ground truth. We depict the results of seven clustering methods that do not require the number of clusters to be known ahead of time.

**Supplementary Figure 2.**
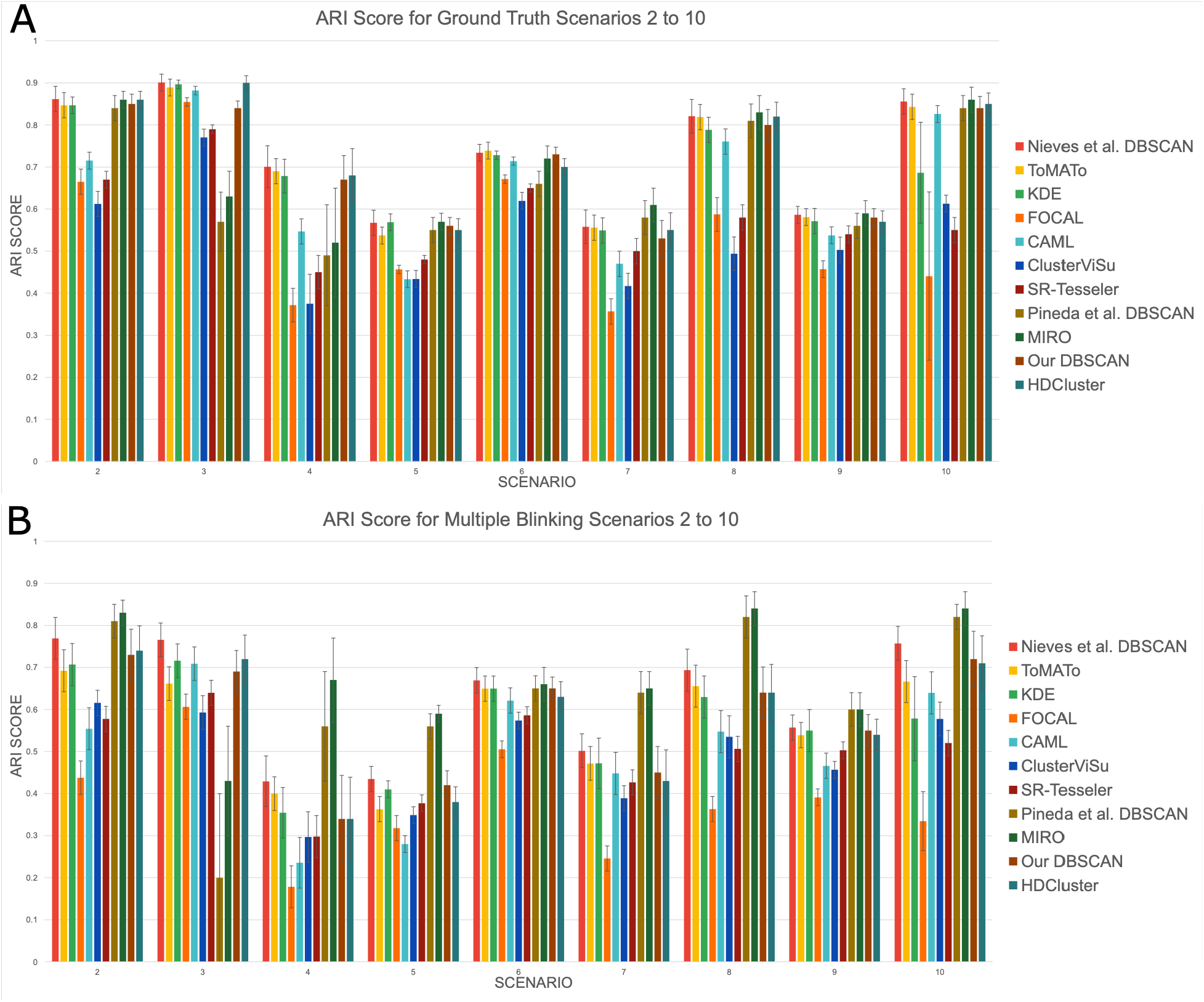
ARI Scores on Scenarios 2–10 of Nieves et al. clustering performance SMLM benchmarking dataset. All methods are compiled from Nieves et al.^26^ results provided in their supplementary material. Scenario 1 was excluded as it was a null hypothesis testing control dataset. (A) Showcases HDCluster performing well with DBSCAN and MIRO as well as other methods on the ideal (i.e., no added blinking) scenarios. (B) The data provide multiple blinking to simulate raw, noisy SMLM data. The figure demonstrates HDCluster performing well alongside our DBSCAN implementation, but lacking by the higher results of DBSCAN as implemented by Nieves et al.^26^. HDCluster still challenges DBSCAN in performance with multiple blinking data.

**Supplementary Figure 3.**
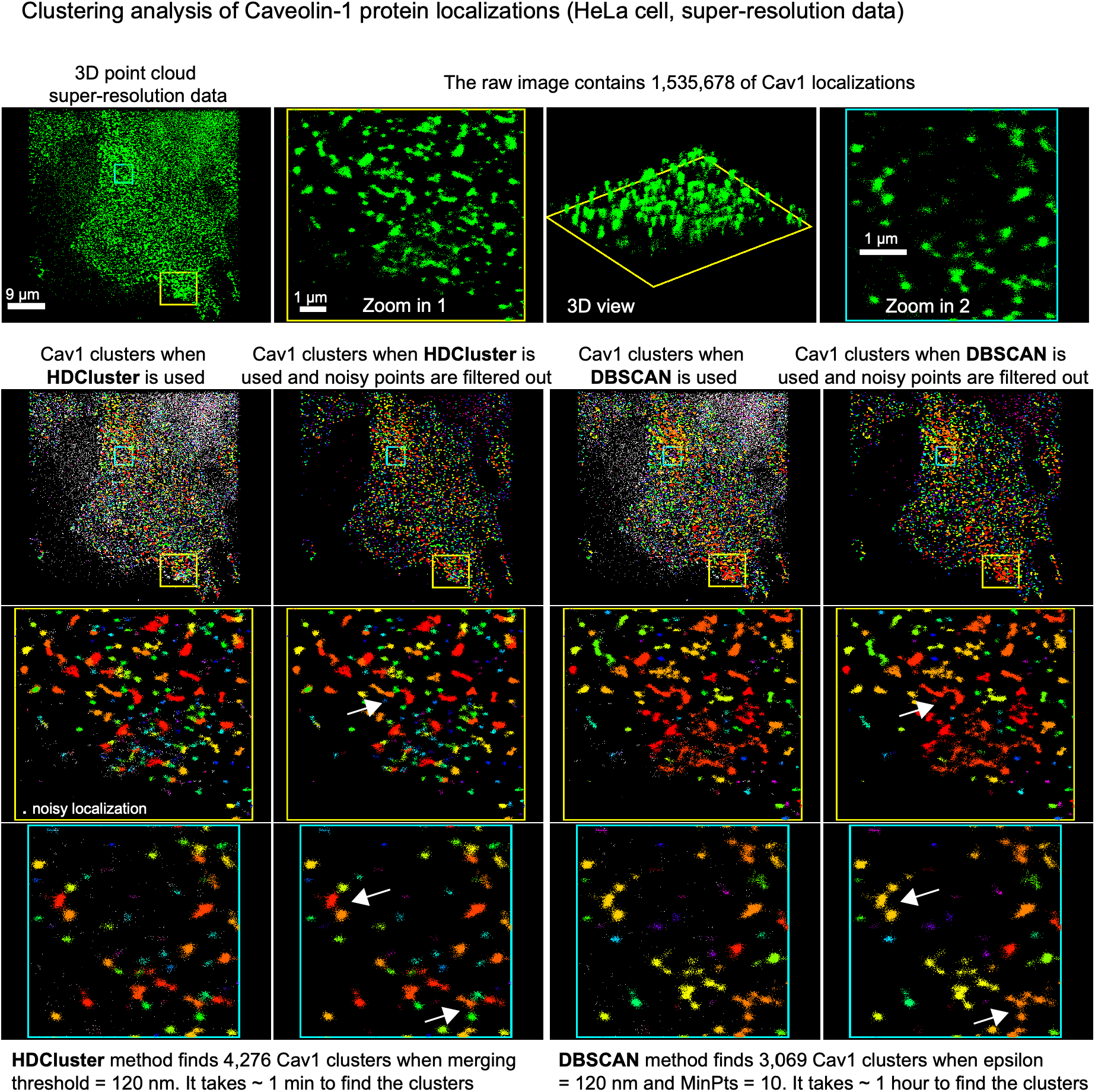
Qualitative analysis comparison of the clustering methods based on the real SMLM dataset. Qualitative analysis showing the results when applying both HDCluster and DBSCAN methods to the Caveolin-1 data of more than 1.5 million localizations^34^. The visualizations show the noisy localizations and the clusters. The arrows in the zoom-in regions show the quality of clustering for the overlapped clusters. The run time analysis for the HDCluster and DBSCAN algorithms on this data is shown in Figure 2E.

**Supplementary Figure 4.**
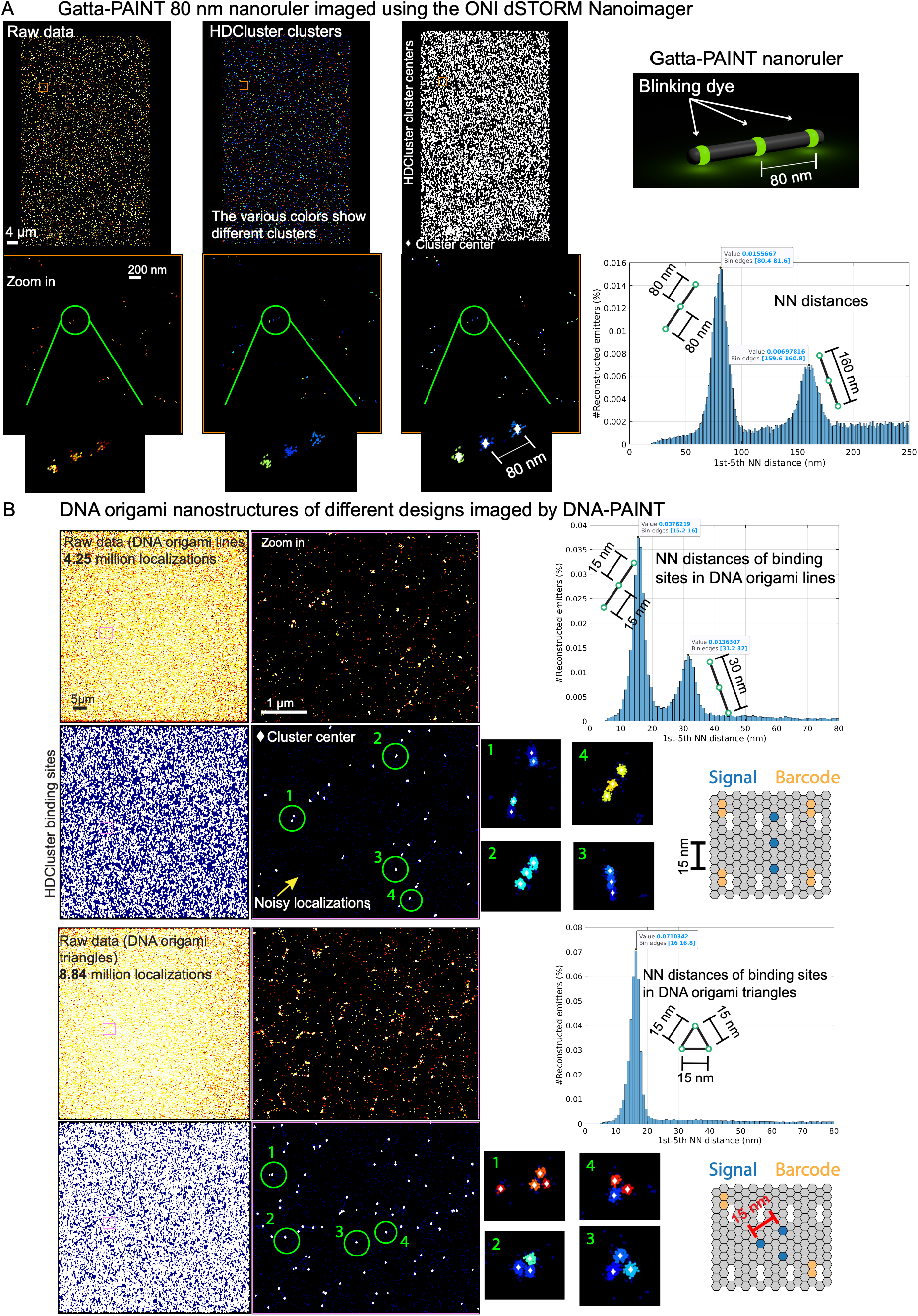
Applying HDCluster to reconstruct the binding sites from dSTORM nanoruler and DNA-PAINT origami nanostructures datasets. (A) Gatta-PAINT 80 nm nanoruler imaged using ONI dSTORM Nanoimager. The HDCluster algorithm is used to denoise, cluster, and reconstruct the emitters of the imaged localizations of the nanorulers. The 1st–5th nearest neighbour (NN) distances of the reconstructed emitters are calculated. The histogram of the 1st–NN distances shows that the majority of reconstructed emitters are distant ∼80 nm from each other, and the second peak of the histogram is at the 2nd–NN of ∼160 nm. (B) DNA-PAINT for two DNA origami nanostructures^38^ is used to illustrate and validate HDCluster denoising, clustering, binding sites reconstruction, and quantification. For the linear nanostructures, HDCluster retrieves the binding sites, and the NN analysis shows the pattern of the binding sites’ stoichiometry. Also, for the triangular nanostructures, HDCluster retrieves the binding sites, and the NN analysis shows the pattern of binding sites stoichiometry that corresponds to the designed origami nanostructures.

**Supplementary Figure 5.**
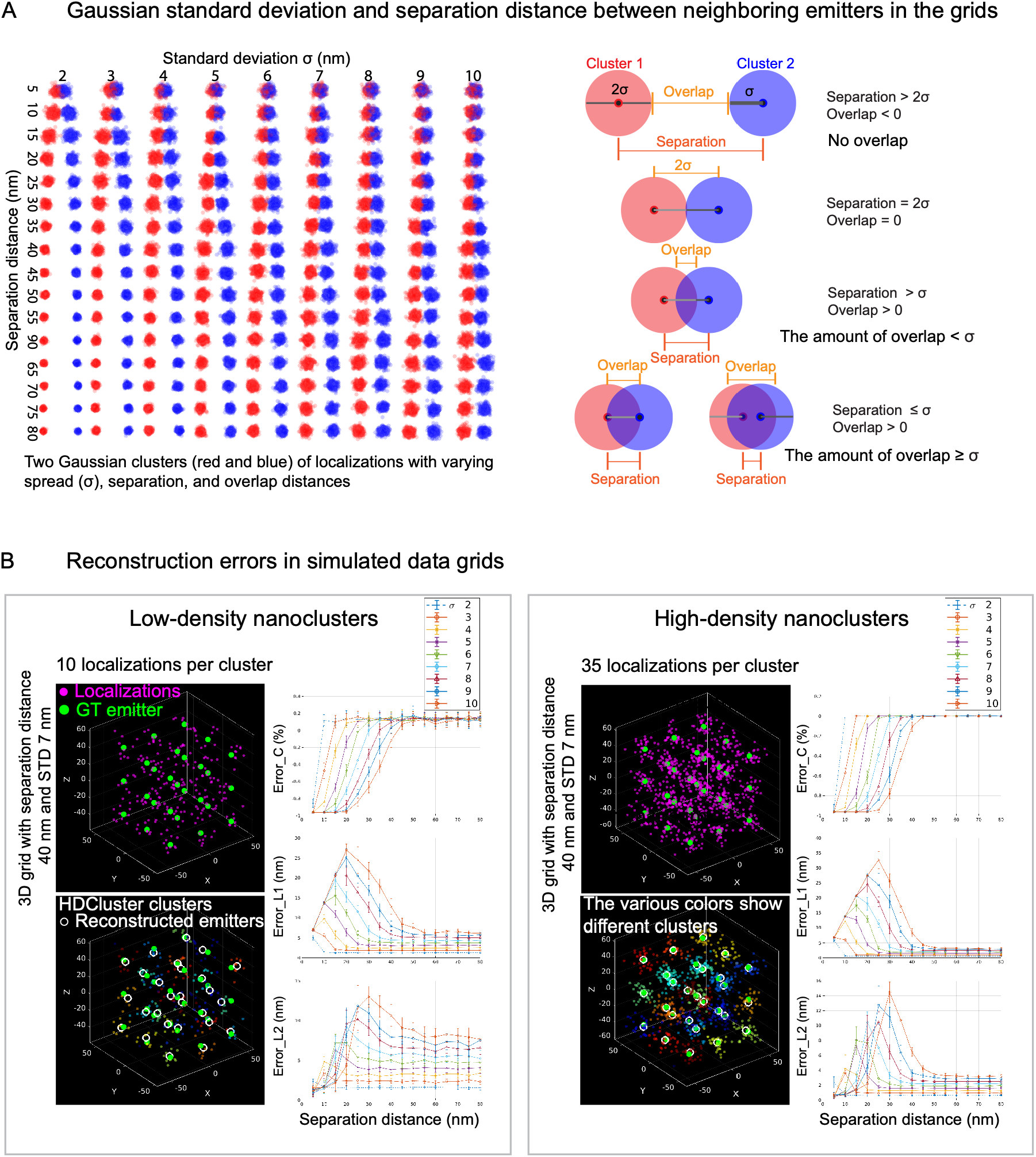
Generation and performance evaluation of synthetic 3D Gaussian clusters for reconstruction error analysis. (A) Generating Gaussian clusters with various standard deviations (σ) and the separation distance values. The cluster’s density (i.e., number of localizations), the spread of localizations around the emitters, and the emitters’ distances can be controlled to assess the performance of the emitters’ reconstruction. σ and separation distance values can be used to set the merging threshold parameter of the HDCluster algorithm (Equation(2)). (B) 20 3D grids of Gaussian clusters with various σ and separation distance values are generated for each pair of σ and separation distance values for a specific cluster density. The counting (*error_C*) and localization errors (*error_L1* and *error_L2*) are calculated for every pair of σ and separation distance values to assess the quality of the reconstruction procedure for emitters/binding sites. Examples for low- and high-density clusters are shown.

**Supplementary Figure 6.**
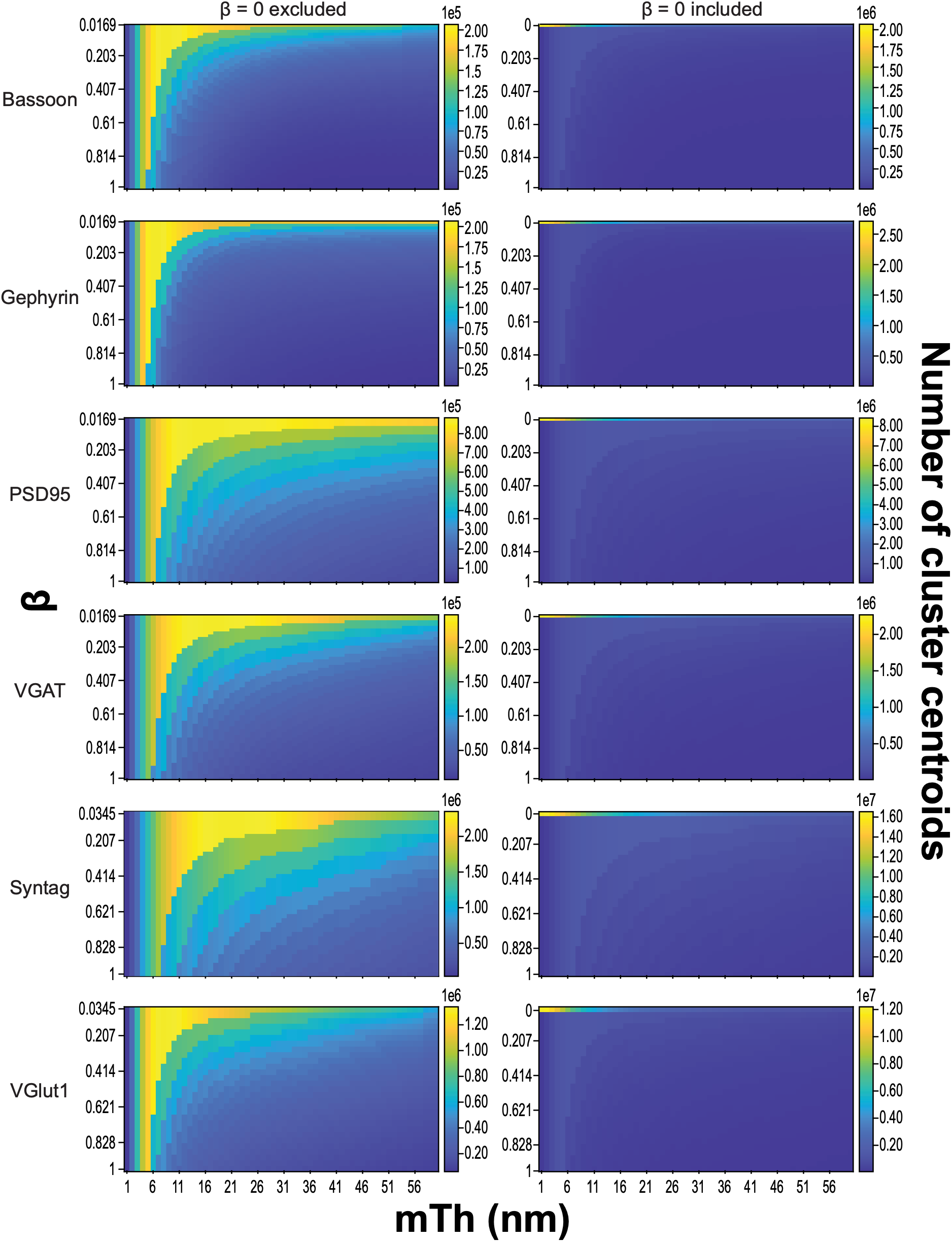
Effect of *mT*ℎ and β on HDCluster cluster-centroid recovery across synaptic-marker channels. Cluster-centroid counts were quantified across combinations of *mT*ℎ from 1 to 60 nm and the optional denoising parameter β from 0 to 1 for the six synaptic-marker channels: Bassoon, Gephyrin, PSD95, VGAT, Synaptotagmin1 and VGlut1. Heatmap color represents the number of cluster centroids recovered for each *mT*ℎ–β combination. Left, β = 0 is excluded to resolve differences among denoising-enabled conditions. Right, the corresponding analysis including β = 0.

