## Supplementary figures and images for "HDCluster: High-Degree Graph Clustering for Robust Analysis of Single Molecule Localization Microscopy"

### Applying HDCluster to reconstruct the binding sites from dSTORM nanoruler and DNA-PAINT origami nanostructures datasets.

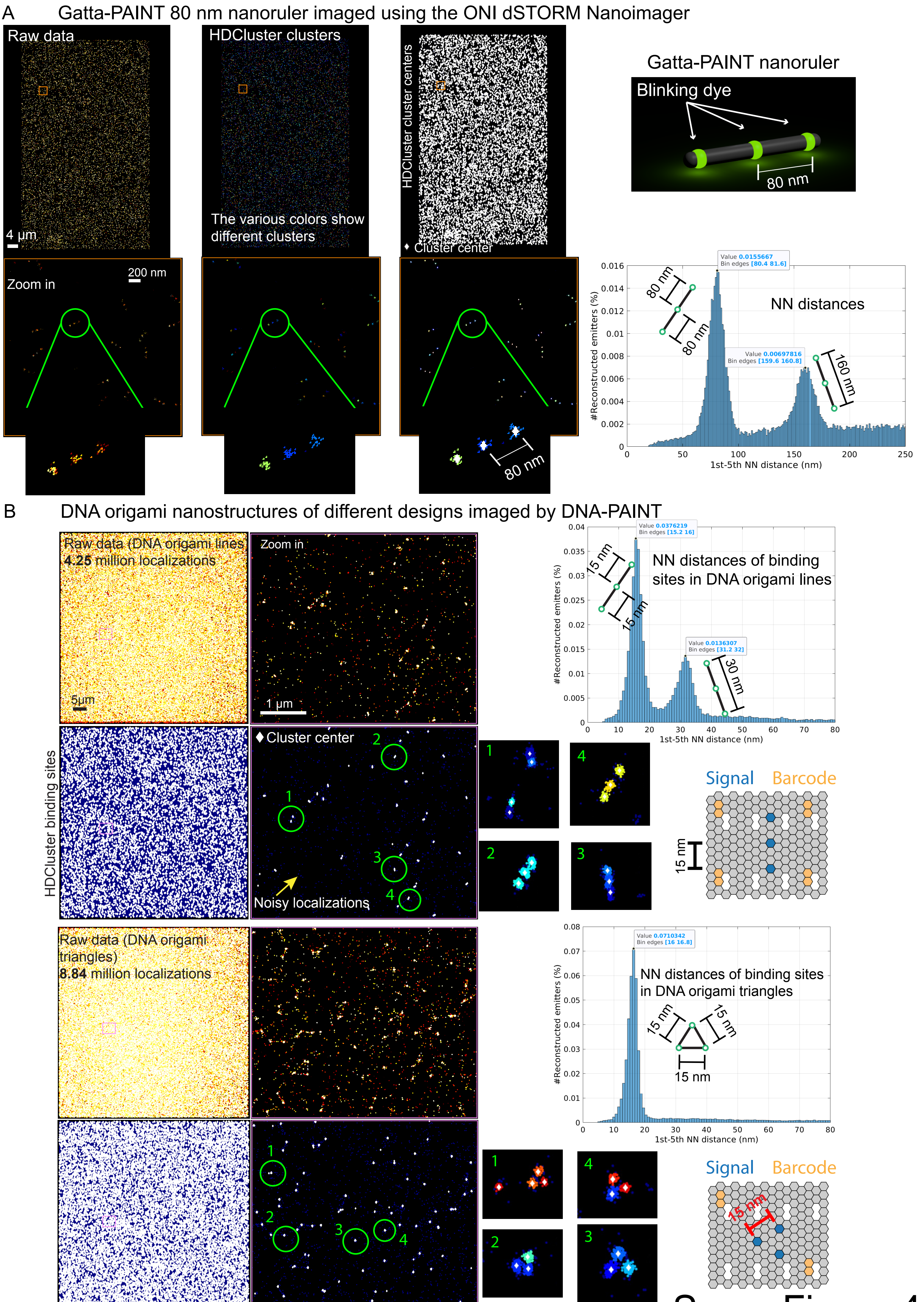


Supp. Figure 4

### Generation and performance evaluation of synthetic 3D Gaussian clusters for reconstruction error analysis.

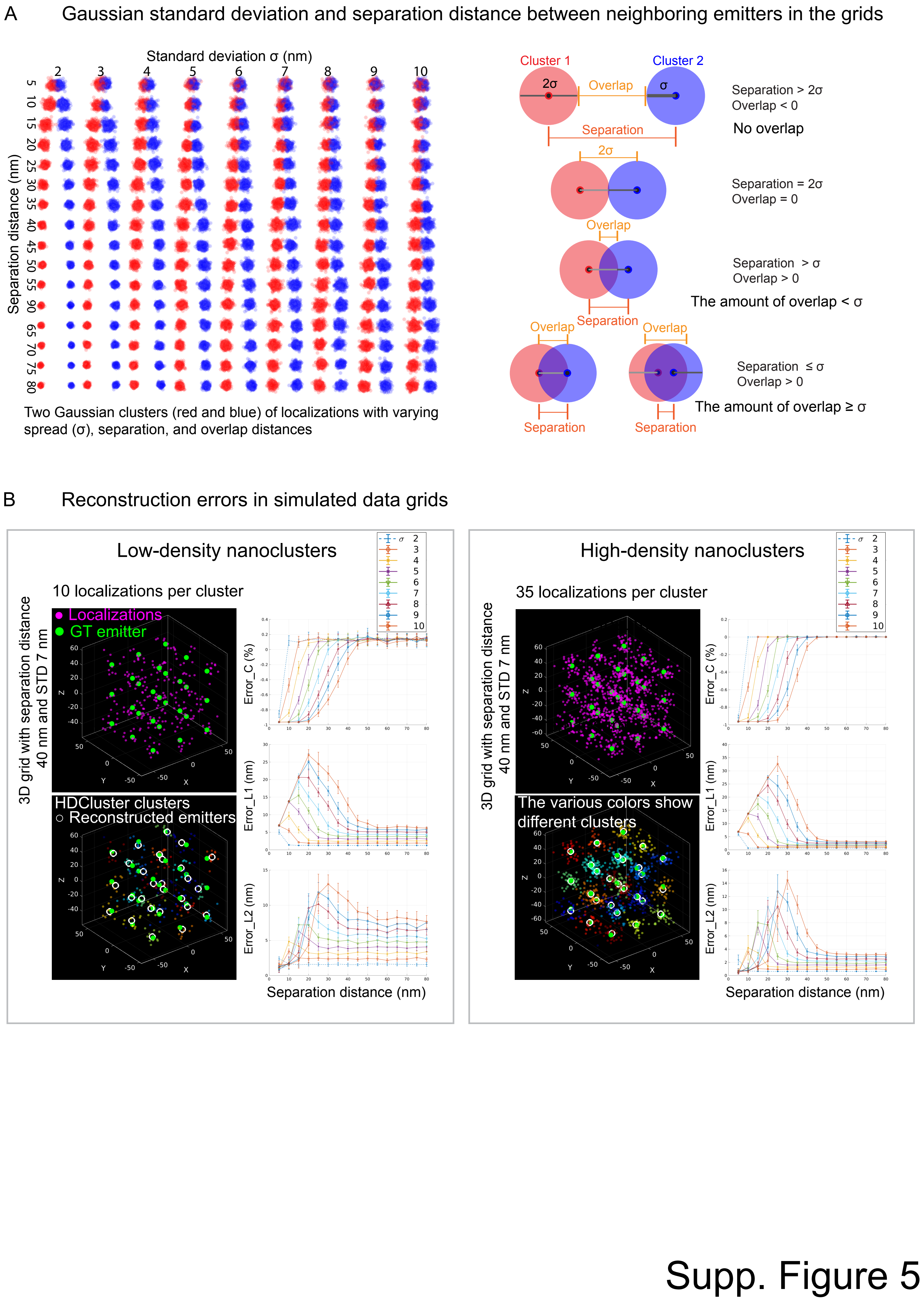

### Supplemental Data 1

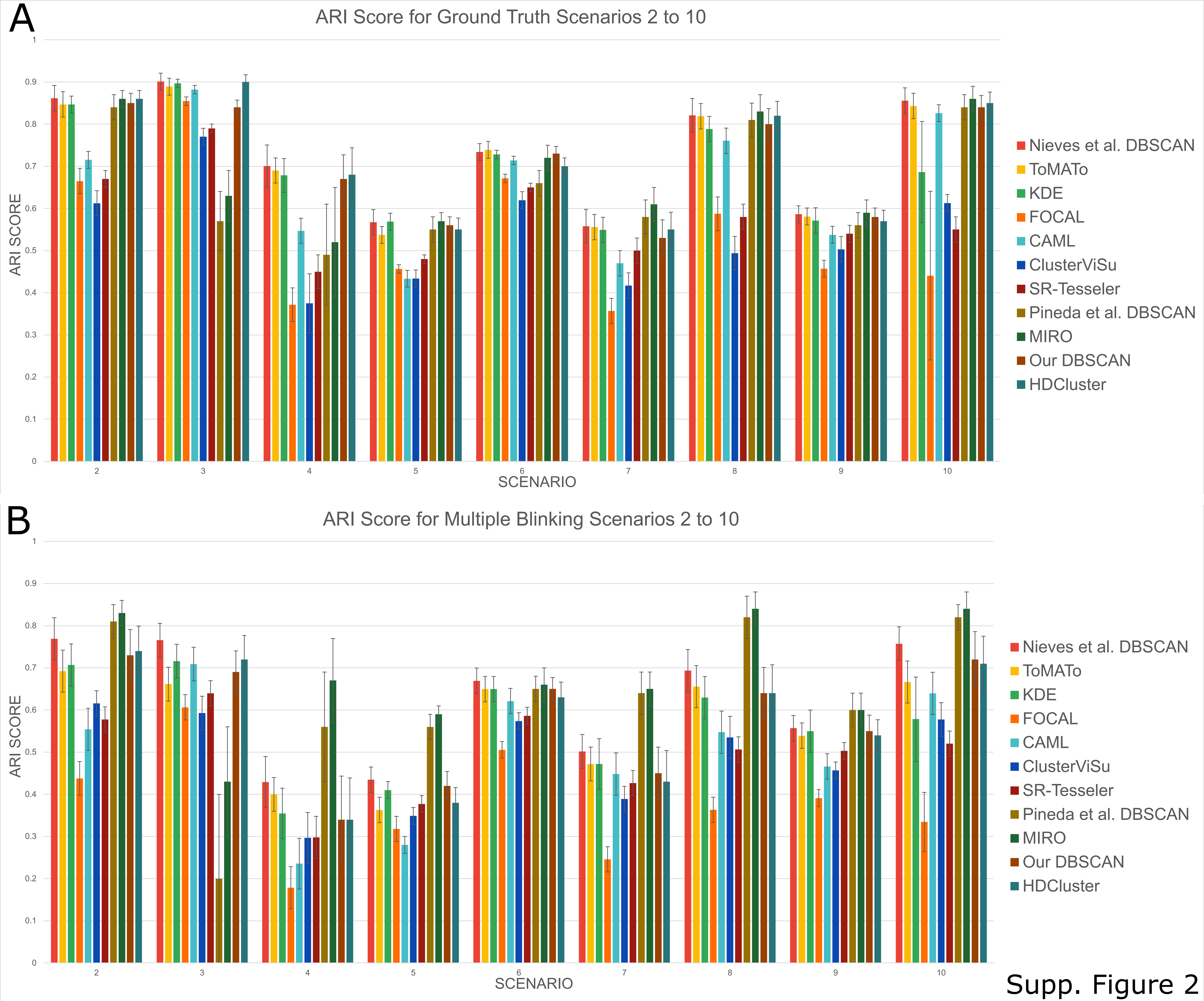


Supp. Figure 2

### Supplemental Data 2

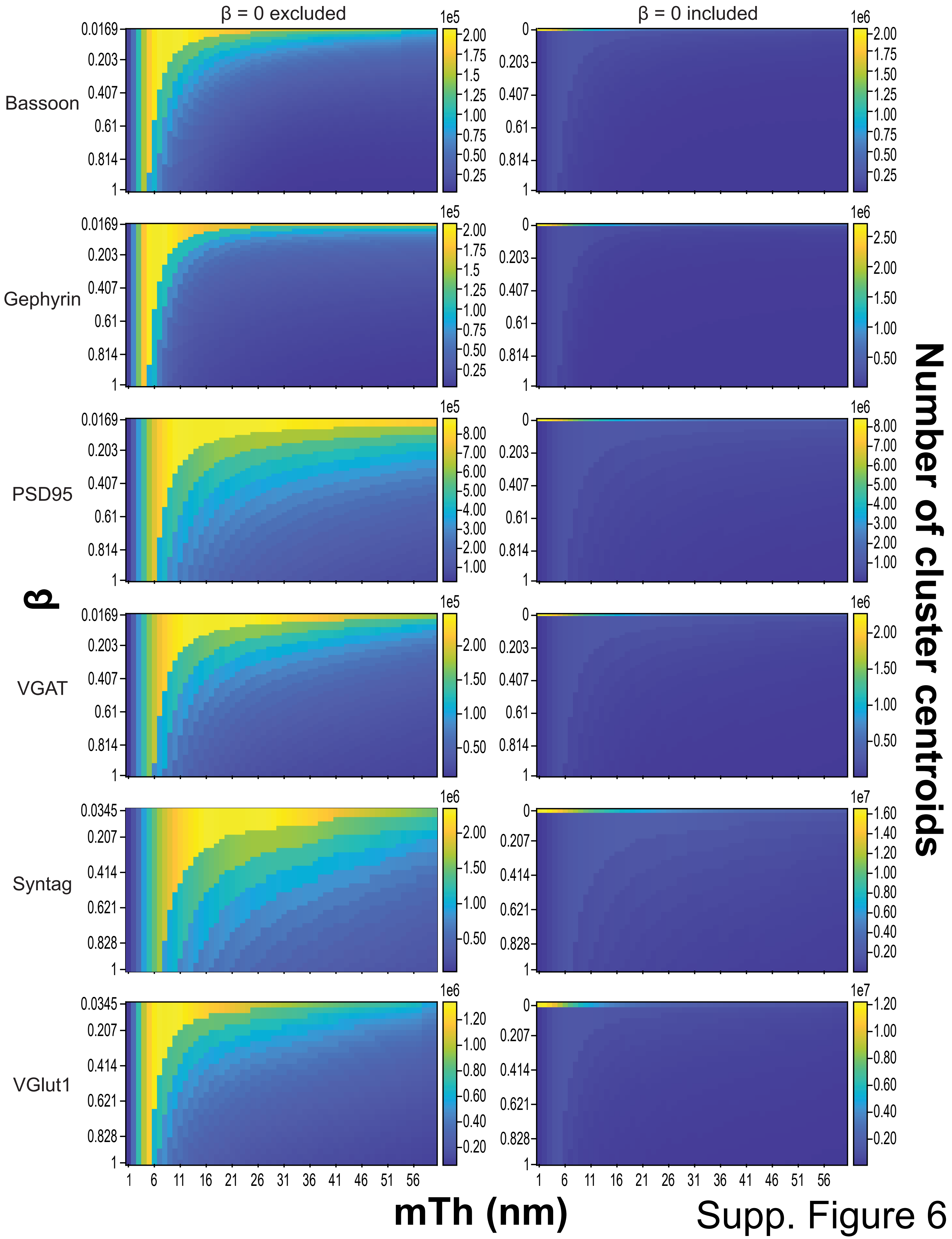


Supp. Figure 6

### Various clustering methods applied to benchmarking shape datasets of a known number of clusters and points' memberships to which cluster.

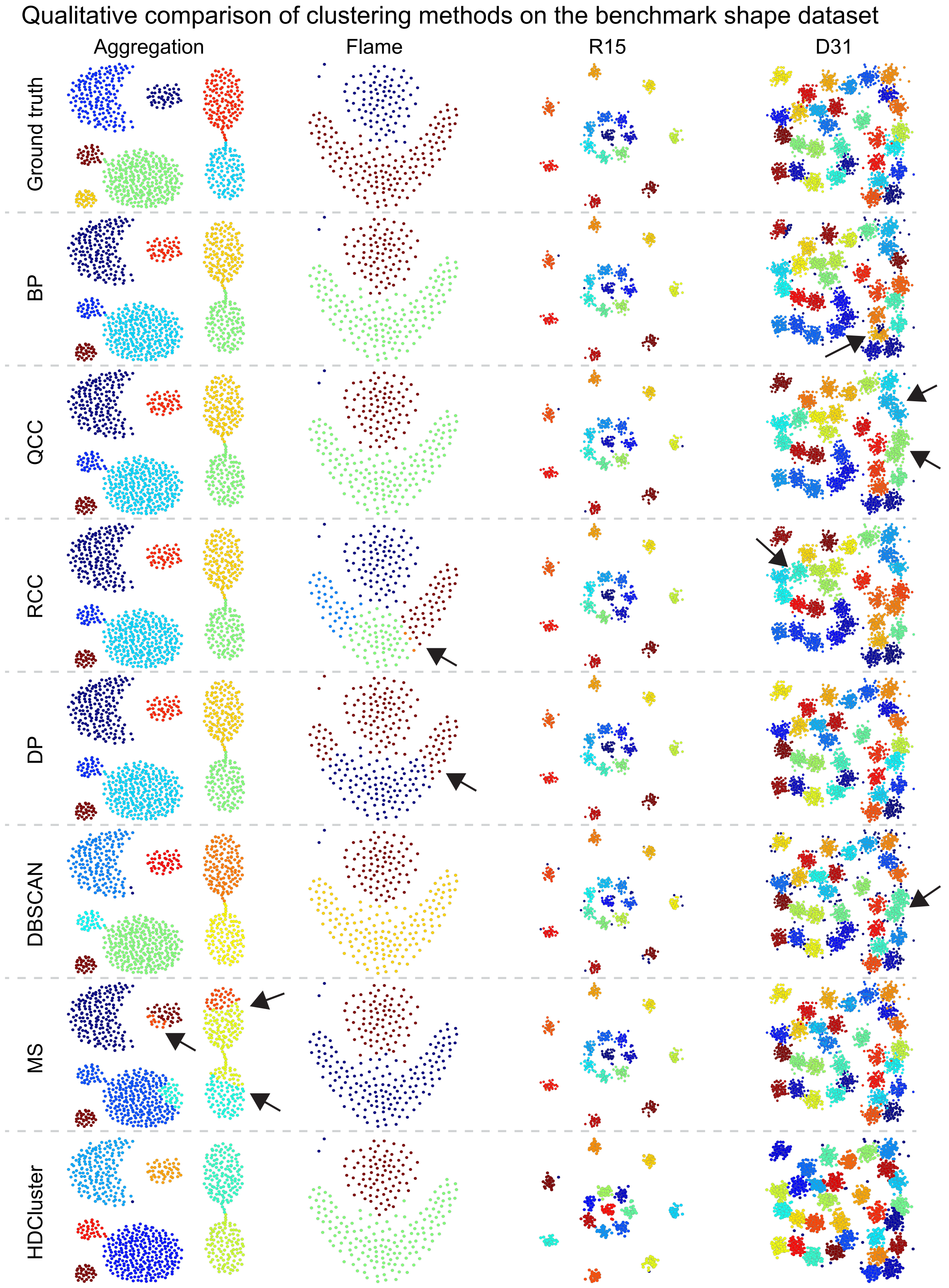


Supp. Figure 1
