## Supplementary material for "HDCluster: High-Degree Graph Clustering for Robust Analysis of Single Molecule Localization Microscopy": Qualitative analysis comparison of the clustering methods based on the real SMLM dataset.

Clustering analysis of Caveolin-1 protein localizations (HeLa cell, super-resolution data)

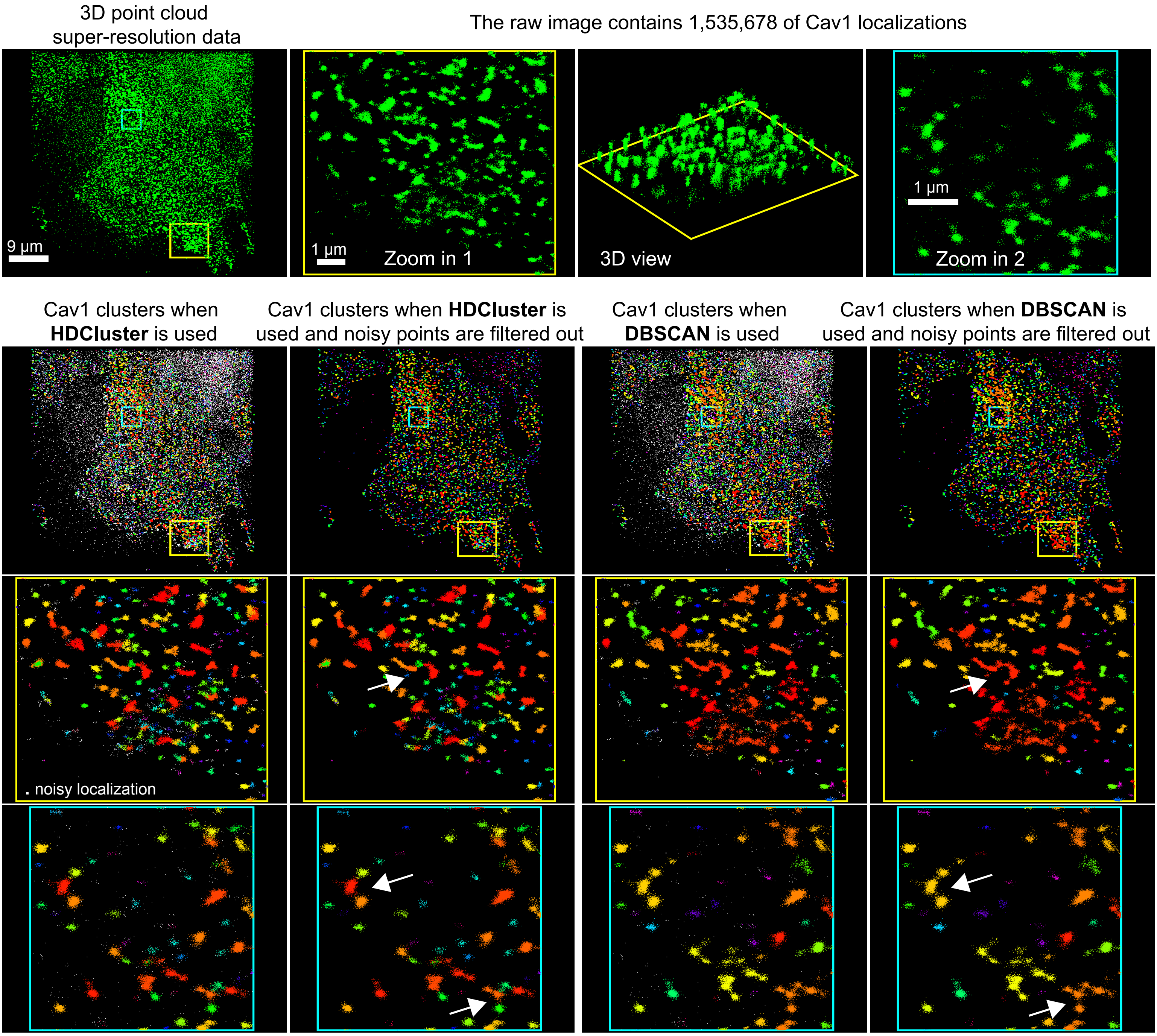


**HDCluster** method finds 4,276 Cav1 clusters when merging threshold = 120 nm. It takes ~ 1 min to find the clusters

**DBSCAN** method finds 3,069 Cav1 clusters when epsilon = 120 nm and MinPts = 10. It takes ~ 1 hour to find the clusters

Supp. Figure 3
